# Strong effect of *Penicillium roqueforti* populations on volatile and metabolic compounds responsible for aromas, flavor and texture in blue cheeses

**DOI:** 10.1101/2020.03.02.974352

**Authors:** Thibault Caron, Mélanie Le Piver, Anne-Claire Péron, Pascale Lieben, René Lavigne, Sammy Brunel, Daniel Roueyre, Michel Place, Pascal Bonnarme, Tatiana Giraud, Antoine Branca, Sophie Landaud, Christophe Chassard

**Affiliations:** Ecologie Systematique Evolution, Université Paris Saclay, CNRS, AgroParisTech, 91400 Orsay, France; Université Clermont Auvergne, INRAE, Vetagro Sup, UMRF, 20 Côte de Reyne, 15000 Aurillac, France; Université Paris-Saclay, INRAE, AgroParisTech, UMR SayFood, 78850 Thiverval-Grignon, France; Laboratoire Interprofessionnel de Production – SAS L.I.P., 34 rue de Salers, 15 000 Aurillac, France

**Author notes:** joint senior authors (Tatiana Giraud, Antoine Branca, Christophe Chassard). Contact information: Thibault Caron.

**Keywords:** Roquefort cheese, fungi, *Penicillium*, domestication, volatile compounds

## Abstract

Studies of food microorganism domestication can provide important insight into adaptation mechanisms and lead to commercial applications. *Penicillium roqueforti* is a fungus with four genetically differentiated populations, two of which were independently domesticated for blue cheese-making, with the other two populations thriving in other environments. Most blue cheeses are made with strains from a single *P. roqueforti* population, whereas Roquefort cheeses are inoculated with strains from a second population. We made blue cheeses in accordance with the production specifications for Roquefort-type cheeses, inoculating each cheese with a single *P. roqueforti* strain, using a total of three strains from each of the four populations. We investigated differences between the cheeses made with the strains from the four *P. roqueforti* populations, in terms of the induced flora, the proportion of blue color, water activity and the identity and abundance of aqueous and organic metabolites as proxies for proteolysis and lipolysis as well as volatile compounds responsible for flavor and aroma. We found that the population-of-origin of the *P. roqueforti* strains used for inoculation had a minor impact on bacterial diversity and no effect on the abundance of the main microorganism. The cheeses produced with *P. roqueforti* strains from cheese populations had a higher percentage of blue area and a higher abundance of the volatile compounds typical of blue cheeses, such as methyl ketones and secondary alcohols. In particular, the Roquefort strains produced higher amounts of these aromatic compounds, partly due to more efficient proteolysis and lipolysis. The Roquefort strains also led to cheeses with a lower water availability, an important feature for preventing spoilage in blue cheeses, which is subject to controls for the sale of Roquefort cheese. The typical appearance and flavors of blue cheeses thus result from human selection on *P. roqueforti,* leading to the acquisition of specific features by the two cheese populations. These findings have important implications for our understanding of adaptation and domestication, and for cheese improvement.

---

Domestication is an evolutionary process that has been studied by many biologists since Darwin. Indeed, domestication is an excellent model for understanding adaptation, because it results from strong and recent selection on traits that are often known and of interest to humans (Larson *et al*., 2014). Furthermore, studies of domestication frequently have important implications for the improvement of cultivated organisms. Domesticated fungi have been less studied than crops in this respect, despite being excellent models in this field (Giraud *et al*., 2017; Gladieux *et al*., 2014). Most fungi can be cultured in Petri dishes, can remain alive for decades when stored in freezers and are propagated asexually. All these features facilitate experimentation. Fungal metabolism results in the production of various compounds of interest, including fuel alcohols, enzymes and antibiotics (Bigelis, 2001). The most ancient and frequent human use of fungi is for fermentation, to preserve and mature food. For example, the yeast *Saccharomyces cerevisiae* is used in the production of bread, wine and beer, and the filamentous fungus *Aspergillus oryzae* is used for the production of soy sauce and sake (Dupont *et al*., 2017). These models have provided important insight into mechanisms of adaptation and domestication (Almeida *et al*., 2014; Baker *et al*., 2015; Gallone *et al*., 2016; Gibbons *et al*., 2012; Gonçalves *et al*., 2016; Libkind *et al*., 2011; Sicard *et al*., 2011).

The *Penicillium* genus contains more than 300 species, several of which are used by humans. For example, penicillin was discovered in *P. rubens,* and two other species, *P. nalgiovense* and *P. salami,* are used in the production of dry-cured meat (Fleming, 1929; Ludemann *et al*., 2010, Perrone *et al*., 2015). For centuries, *Penicillium roqueforti* (Thom) has been used in the maturation of all blue cheeses worldwide (Coton *et al*., 2020a; Labbe *et al*., 2004, 2009; Vabre, 2015). This fungus generates cheeses with a blue-veined appearance, by producing melanized conidia within cavities in the cheese in which oxygen is readily available (Moreau, 1980). *Penicillium roqueforti* is also found in non-cheese environments (Pitt *et al*., 2009; Ropars *et al*., 2012), and four genetically differentiated clusters of individuals (i.e., populations) have been identified in *P. roqueforti*. Two populations are used for cheesemaking, whereas the other two populations thrive in silage, lumber or spoiled food (Dumas *et al*., 2020; Gillot *et al*., 2015; Ropars *et al*., 2014). Genomic and experimental approaches have provided compelling evidence for the domestication of cheese *P. roqueforti* populations (Cheeseman *et al*., 2014; Dumas *et al*., 2020; Gillot *et al*., 2015, 2017; Ropars *et al*., 2015, 2016b, 2017) which have been reviewed (Coton *et al*., 2020b; Ropars *et al*., 2020a). Indeed, the populations of *P. roqueforti* used to make blue cheeses display the characteristic features of domesticated organisms: genetic and phenotypic differences relative to non-cheese populations, with, in particular, traits beneficial for cheese production, such as faster growth on cheese medium (Dumas *et al*., 2020; Gillot *et al*., 2015; Ropars *et al*., 2014, 2016), but also lower fertility and lower fitness in nutrient-poor conditions (Ropars *et al*., 2015, 2016). Both cheese populations have lower levels of genetic diversity than the two non-cheese populations, indicating an occurrence of bottlenecks (Dumas *et al*., 2020), which typically occur during domestication. The two cheese populations are genetically and phenotypically differentiated from each other, suggesting that they result from independent domestication events (Dumas *et al*., 2020). One of the cheese populations, the non-Roquefort population, is a clonal lineage with a very low level of genetic diversity used to produce most types of blue cheeses worldwide. The second cheese population, the Roquefort population, is genetically more diverse and contains all the strains used to produce blue cheeses from the emblematic Roquefort protected designation of origin (PDO) (Dumas *et al*., 2020). *In vitro* tests have shown that the non-Roquefort population displays faster tributyrin degradation (*i.e.* a certain type of lipolysis) and higher salt tolerance, faster *in vitro* growth on cheese medium, better exclusion of competitors than the Roquefort population (Dumas *et al*., 2020; Ropars *et al*., 2014, 2015), and an absence of toxic mycophenolic acid production (Gillot *et al*., 2017). The specific features of the Roquefort population may result from the constraints of the PDO, requiring the use of local strains and at least 90 days of maturation, and preventing the use of strains from the non-Roquefort population better suited to modern modes of production (Dumas *et al*., 2020). Genomic footprints of domestication (i.e., of adaptive genetic changes) have been identified in the two *P. roqueforti* populations used for cheesemaking. Indeed, it has been suggested that horizontally transferred genes found only in the non-Roquefort population are involved in the production of an antifungal peptide and in lactose catabolism (Cheeseman *et al*., 2014; Ropars *et al*., 2014, 2015). The effects of positive selection have been detected in genes with predicted functions in flavor compound production, in each of the cheese populations (Dumas *et al*., 2020).

Thus, the four *P. roqueforti* populations probably harbor multiple specific traits, leading to the generation of cheeses with different physicochemical properties and flavors, although this has yet to be tested. Assessments of the effect of the population-of-origin of the *P. roqueforti* strain used on the features of the cheese will i) provide important fundamental knowledge about the trait under selection for cheesemaking and adaptation to the cheese environment, ii) provide a basis for the elucidation of other genomic changes and iii) be crucial to improvements in strain use and improvement. *Penicillium roqueforti* is used as a secondary starter for flavor production, mostly through proteolysis (*i.e.* casein catabolism) and lipolysis during ripening (Moreau, 1980). The main characteristic feature of blue cheeses, and of Roquefort PDO cheeses in particular, is their intense, spicy flavors (Kinsella *et al*., 1976; Rothe *et al*., 1982). The specific volatile and metabolic compounds responsible for these flavors are generated principally by lipolysis in blue cheeses (Cerning *et al*., 1987; Collins *et al*., 2003) and their intensity varies among *P. roqueforti* strains (Dumas *et al*., 2020; Gillot *et al*., 2017; Larsen *et al*., 1999). The fatty acids released by lipolysis are the precursors of aldehydes, alcohols, acids, lactones and methyl ketones, which provide the moldy aromas typical of blue cheeses (Collins *et al*., 2003). *Penicillium roqueforti* degrades most proteins, but the efficiency of proteolysis differs between strains (Cerning *et al*., 1987; Dumas *et al*., 2020; Gillot *et al*., 2017; Larsen *et al*., 1998). The resulting peptides contribute to flavors, and their degradation into amino acids further influences cheese aroma and the growth of other microorganisms (McSweeney *et al*., 2000; Williams *et al*., 2004). *Penicillium roqueforti* also contributes to lactate degradation, which is necessary for deacidification and promotes the development of less acid-tolerant microorganisms (McSweeney *et al*., 2017). Through these effects, and by producing secondary metabolites with antimicrobial properties, *P. roqueforti* may also affect the microbial composition of the cheese (Kopp *et al*., 1979; Vallone *et al*., 2014). Another parameter potentially affected by *P. roqueforti* populations and restricting the occurrence of spoiler microorganisms is the lack of free water, (i.e., low water activity, Aw), which is subject to strong controls for the sale of Roquefort cheese and is affected by the degree of proteolysis (Ardö *et al*., 2017). The *P. roqueforti* population may, thus, also have an indirect effect on the features of the cheese, through various effects on beneficial or undesirable contaminants.

The differences between *P. roqueforti* populations have, to date, been studied only *in vitro* or in very rudimentary cheese models. Here, our objective was to assess the effect of the *P. roqueforti* population-of-origin of the strain used for inoculation on the features of blue cheeses. We focused on several features considered important for cheese quality. Given the evidence from previous studies that cheese *P. roqueforti* populations have been domesticated, any differences between the cheeses produced with cheese and non-cheese populations, and/or between the two cheese populations can probably be considered to reflect selection by humans for the production of good cheeses, either on standing variation in the ancestral *P. roqueforti* population or for *de novo* mutations. We therefore produced blue cheeses in conditions very similar to those used in industrial Roquefort PDO production, using, in particular, milk from the local “Lacaune” breed, with strains from the four *P. roqueforti* populations. We compared several important cheese features between the four populations: i) physicochemical features, relating to texture and biochemical composition, ii) cheese microbiota composition and abundance, which may have effects on several cheese features, iii) the blue area as a proportion of cheese slices, which is important for the blue-veined appearance of the cheese and is dependent on the growth and sporulation of *P. roqueforti* in cheese cavities, and iv) the types and abundance of the metabolic and volatile compounds produced, which influence flavor and aroma. We investigated the differences in these features between the cheeses produced with strains from the four *P. roqueforti* populations (Roquefort cheese, non-Roquefort cheese, silage and lumber/food spoiler populations). We also investigated the possible differences between cheeses made with cheese and non-cheese populations, and between the Roquefort and non-Roquefort cheese populations. Assessments of the traits differing between cheese and non-cheese *P. roqueforti* populations, and between the two cheese populations, and investigations of whether the cheese populations are more suitable for cheese-making, are of fundamental importance for understanding the domestication of cheese fungi, through the identification of traits subjected to selection. Furthermore, the identification of differences between *P. roqueforti* populations of relevance for cheese making has many applied consequences for the cheese industry, in terms of strain choice for different kinds of blue cheeses, paving the way for the improvement of mold strains by generating progenies from crosses of the two cheese populations, and for the choice of traits for measurement and selection in offspring.

## Materials and Methods

The Materials and Methods are described in more details in the Supplementary Methods.

### Experimental design and fungal strains

We made cheeses with three different strains from each of the four *P. roqueforti* populations (Figure 1, a strain is defined here as a haploid individual obtained by monospore isolation), according to population assignments obtained in a previous study based on molecular markers (Dumas *et al*., 2020). For cheese production (Figure 1B), we used a single strain per cheese for inoculation, using a total of three different strains from each of the four *P. roqueforti* populations (Figure 1A). Due to the limited production capacity of the experimental facility, we were unable to make all the cheeses at the same time. We therefore split production into three assays, each including one strain from each of the four populations (Figure 1A). For each strain in each assay, we created three production replicates, with two cheeses per strain in each replicate, to ensure the production of sufficient material for sampling. In total, we produced 72 cheeses (4 strains * 2 cheeses * 3 replicates * 3 assays, Figure 1A). The assays were performed sequentially from February to April. The effect of the seasonal change in milk composition was therefore confounded with the strain effect within the population, hereafter referred to as the “assay effect”. The three replicates within each assay were also set up at different times, a few days apart, and thus with different batches of raw milk (Figure 1A). LCP (*Laboratoire de Cryptogamie de Paris*) strains were obtained from the *Muséum d’Histoire Naturelle de Paris*, France, and BRFM (Biological Resource Fungi Marseille’) strains were obtained from the *Centre International de Ressources Microbiennes de Marseille*, France.

**Figure 1:**
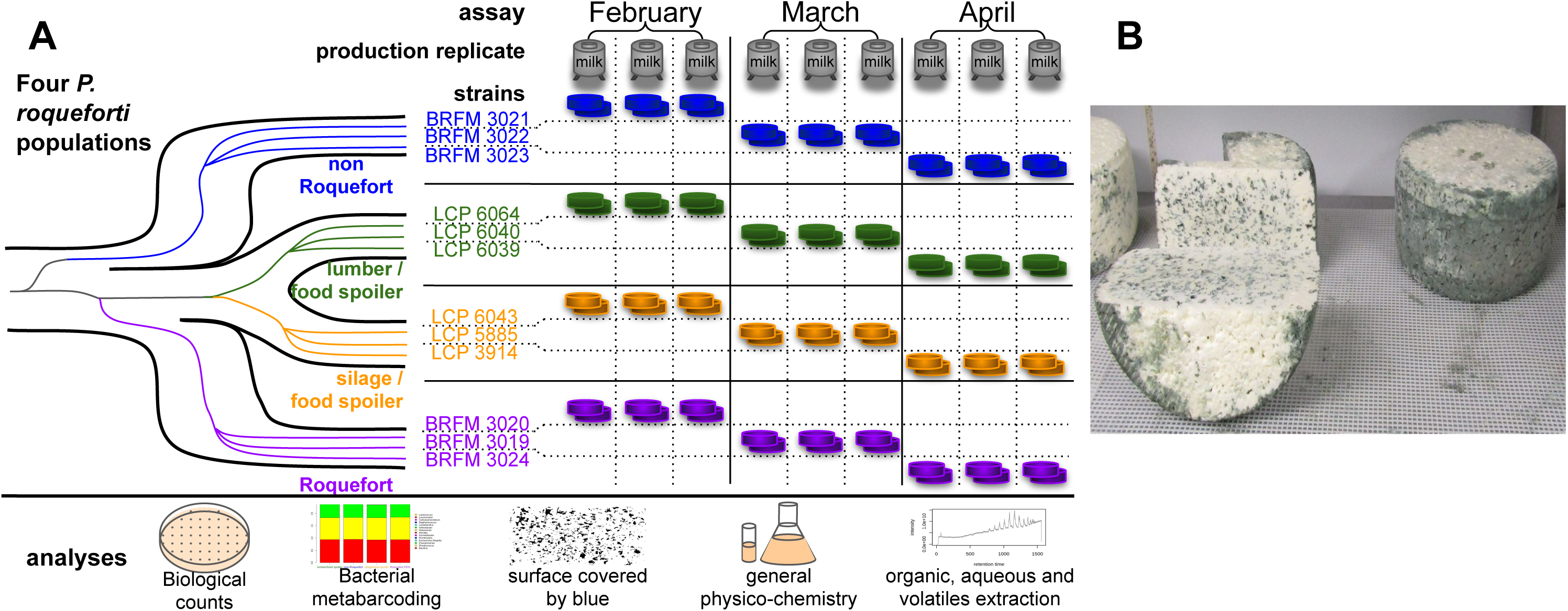
Experimental cheesemaking. (A) Experimental design for cheesemaking, using one strain per cheese, and three different strains from each of the four *Penicillium roqueforti* populations (non-Roquefort cheese in blue, Roquefort cheese in purple, silage/food spoiler in orange, lumber/food spoiler in green, the lineages of which are shown on the left). Each assay (February, March, April) corresponded to a single strain from each of the four populations, with three production replicates at different times, different batches of unpasteurized milk and with two cheeses produced per strain in each replicate. The identities of the strains used are indicated on the left of each assay, for each of the four *P. roqueforti* populations. (B) Photograph of the experimental cheeses after 20 days of maturation.

### Cheesemaking

The cheesemaking protocol was typical of the procedures used by the main producers of Roquefort cheese and complied with the Roquefort PDO specifications, except that the ripening process took place in artificial cellars in the INRA facilities at Aurillac and with strains from different *P. roqueforti* populations. Briefly, we slowly mixed about 35 L of milk in each vat, and heated it to 32.5°C. We then added 20 mg.L^-1^ mesophilic starter culture containing *Leuconostoc* spp., *Lactococcus lactis* subsp. *cremoris*, *L. lactis* subsp. *lactis*, and *L. lactis* subsp. *lactis* biovar *diacetylactis* (FD-DVS CHN-11, CHR HANSEN, Saint-Germain-lès-Arpajon, France), 25 mg.L^-1^ gas-producing *Leuconostoc mesenteroïdes* subsp. *mesenteroïdes* (CHOOZIT LM 57 LYO 20 DCU, DANISCO, Paris, France) and about 3.6 x 10^6^ CFU.L^-1^ *P. roqueforti* spores (SAS LIP, Aurillac, France). The mixture was homogenized for five minutes, and we then added 0.25 ml of active chymosin at a concentration of 520 mg.L^-1^ (Laboratoires Humeau, La Chapelle-sur-Erdre, France). The resulting curd was cut into 1.5 cm^3^ cubes, left to rest for five minutes, and allowed to ferment for 50 minutes. The curd was drained and placed in perforated cylinders of 20×10 cm at 26°C/85% humidity. The cheeses were turned five times after molding. On the second day, the cheeses were unmolded. On the third day, the cheeses were rubbed with sterilized coarse salt and transferred to ripening cellars at 11°C / 95% humidity. On the fifth day, the cheeses were resalted in the same way. On the seventh day, the cheeses were pricked. They were left to ripen until day 20, when they were wrapped in sterile aluminum foil and allowed to mature at -2°C until day 180.

Cheese samples were collected on days 0, 9, 20, 90 and 180. These time points are referred to hereafter as “stages”. Day 0 corresponded to raw milk for microbiology analysis and sowed curd for metabolic and volatile compound analyses. Days 9 and 20 corresponded to half-way through and the end of the ripening period, respectively. Days 90 and 180 (i.e. 3 and 6 months) corresponded to the minimum maturation times for the Roquefort PDO and a typical maturation period for sale, respectively.

### Microbial and metabarcoding analyses

We estimated the concentrations of various microbial communities in the initial unpasteurized milk and at various stages of cheese maturation, to determine whether the *P. roqueforti* population affected the cheese microbiota (for more information, see the Supplementary Methods). We counted the total number of aerobic mesophilic bacteria on plate count agar (PCA; Nelson, 1940). Mesophilic lactic acid bacteria were counted on Man Rogosa Sharpe agar (MRS; De Man *et al*., 1960) and thermophilic lactic acid bacteria were counted on M17 agar (Terzaghi, 1975). Dextran-positive *Leuconostoc* spp. were counted on Mayeux, Sandine and Elliker medium (MSE; Mayeux *et al*., 1962). Molds and yeasts were counted on oxytetracycline gelose agar medium (OGA; Mossel *et al*., 1970). Gram-positive catalase-positive bacteria were counted on cheese-ripening bacterial medium (CRBM; Denis *et al*., 2001). Enterobacteria were counted on violet red bile glucose (VRBG; Mossel *et al*., 1978). We further investigated the possible effects of the *P. roqueforti* population on the identity and relative abundance of other microorganisms in cheeses, by performing metabarcoding analyses on our experimental cheeses at 9 and 20 days (during the ripening period), by extracting DNA, and amplifying and sequencing the V3-V4 region of the 16S rDNA gene. Amplicons were sequenced with Illumina Miseq technology. The mean read depth of the 71 samples (36 at 9 and 20 days of ripening minus one lost) was 28539 reads. Amplicon data from high-throughput sequencing were analyzed with Find Rapidly OTUs in Galaxy Solution (FROGS) v3.0 (Escudié *et al*., 2018). For each OTU, taxonomic assignment was determined with the Silva-132 (https://www.arb-silva.de/) and 16S rDNA RefSeq databases (https://blast.ncbi.nlm.nih.gov/Blast.cgi). Four widely used diversity parameters were calculated from OTU compositions: two for alpha diversity (Shannon index, Simpson index) and the Bray-Curtis dissimilarity for beta diversity.

### Blue area

We estimated the percentage area of the cheese that was blue, on fresh inner cheese slices (Figure 3B). This percentage is dependent on the formation of cavities within the cheese, the growth of *P. roqueforti* within the cavities and its sporulation, as the blue color is due to the melanin present in *P. roqueforti* spores. We cut all the 20-, 90- and 180- day cheeses in half and took three photographs of each fresh slice with a Canon PowerShot SX410 IS (JPEG format, 5152 x 3864 pixels, 100 ISO, without flash). We analyzed these images with imageJ 1.52n (Schneider *et al*., 2012; Figure S2): (i) the brightness and contrast of the raw images were standardized with a stack contrast adjustment plugin, using a reference image (Figure S2A,B) (ii) rectangular selection was used to crop the images manually to ensure that they contained only the cheese slices (Figure S2C), (iii) the red channel was segmented with a grayscale threshold of 102 to distinguish colonized cavities from the inner white parts of the cheese and empty cavities (Figure S2D), (iv) the number of dark pixels in the cavities was counted and divided by the total number of pixels for the entire cheese slice, to determine the percentage blue area of the slice. Within the dark patches in the cheese cavities colonized by the fungus, all pixels were considered dark (script at https://gitlab.com/snippets/1945218).

### Physicochemistry

We performed standard physicochemical measurements on the cheeses. We measured dry matter content, fat content as a proportion of dry matter, the moisture content of the defatted cheese, total, soluble and non-protein nitrogen contents, chloride and salt content, water activity and pH at various stages of maturation, according to reference methods (for more information, see the Supplementary Methods). We measured glucose, lactose, lactate, acetate and butyrate concentrations in the cheeses on days 9 and 20, by high-performance liquid chromatography (HPLC, for more information, see the Supplementary Methods).

### Metabolic and volatile compounds

We investigated possible differences in proteolytic and lipolytic activities between the four populations, by UHPLC-MS after two extraction procedures (in water and an organic solvent). We analyzed the abundance of free fatty acids and residual glycerides in 90-day cheeses, by coupling a global extraction (accelerated solvent extraction with hexane-isobutanol) with UHPLC-MS analysis in the positive (triglycerides) and negative (fatty acids) ionization modes (for more information, see the Supplementary Methods). We investigated the identity and abundance of volatile flavor and aroma compounds, using a dynamic headspace system (DHS) with a Gerstel MPS autosampler (Mülheim an der Ruhr, Germany) and gas chromatography-mass spectrometry analysis with an Agilent 7890B GC system coupled to an Agilent 5977B quadrupole mass spectrometer (Santa Clara, United States). Statistical analyses were performed with R software (http://www.r-project.org/). Further details about the materials and methods are provided in the Supplementary Methods.

## Results

### *Penicillium roqueforti* population-of-origin influences bacterial diversity slightly, but has no effect on the abundance of the main microorganisms in the cheese

We investigated whether the population-of-origin of the *P. roqueforti* strains used for inoculation affected the composition of the cheese microbiota, by estimating the densities of key microbial communities through colony counts (CFU/g) on various specific culture media and a metabarcoding approach based on 16S sequencing targeting the bacteria in cheeses at several stages of maturation. Based on microbial counts on plate, we found no significant effect of *P. roqueforti* population on the abundance of any of the counted microorganisms, including molds (i.e. *P. roqueforti)*, at any sampled stage of maturation (Supplementary Figure 1; Supplementary Table 1A).

The metabarcoding approach targeting bacteria identified mostly sequences from the *Lactococcus* and *Leuconostoc* spp. starters (about 58 and 40%, respectively, Figure 2A, Supplementary Table 1B), which are responsible for acidification and cavity formation, respectively, in the cheese. The remaining sequences corresponded to 12 bacterial genera frequently found in raw milk cheeses, such as *Lactobacillus*, *Staphylococcus* and *Arthrobacter* (<1%, Supplementary Table 1B). However, the large predominance of starters made it impossible to obtain sufficient data for other bacteria to assess differences in the abundance of particular bacteria between cheeses made with strains from the four *P. roqueforti* populations (Supplementary Table 1C). We also targeted eukaryotes, based on ITS sequences, in a few samples. Most sequences were assigned to *P. roqueforti* spp. (89%) and, to a lesser extent, *Candida* spp. (10%, data not shown), so no further analyses were performed on the remaining low-abundance sequences. We estimated three OTU (operational taxonomic unit) diversity parameters based on bacterial barcode sequence abundances, to measure OTU richness and/or evenness. Bray-Curtis dissimilarity showed that cheeses made with strains from the same *P. roqueforti* population were no more similar than those made with strains from different *P. roqueforti* populations. However, we found a significant effect of *P. roqueforti* population, in addition to a stage effect, on the Shannon and Simpson diversity indices (Figures 2B and 2C). Cheeses made with strains from the cheese *P. roqueforti* populations tended to have a higher bacterial OTU diversity, particularly at nine days of maturation and for the Roquefort population (Figures 2B and 2C), although the post-hoc analyses were not powerful enough to detect significant pairwise differences (Supplementary Table 1C). There may also be undetected differences at species level or for low-abundance microorganisms that might nevertheless have substantial effects. However, even if this were the case, it would constitute an indirect effect of the *P. roqueforti* population, as this was the only difference during our cheesemaking process.

**Figure 2:**
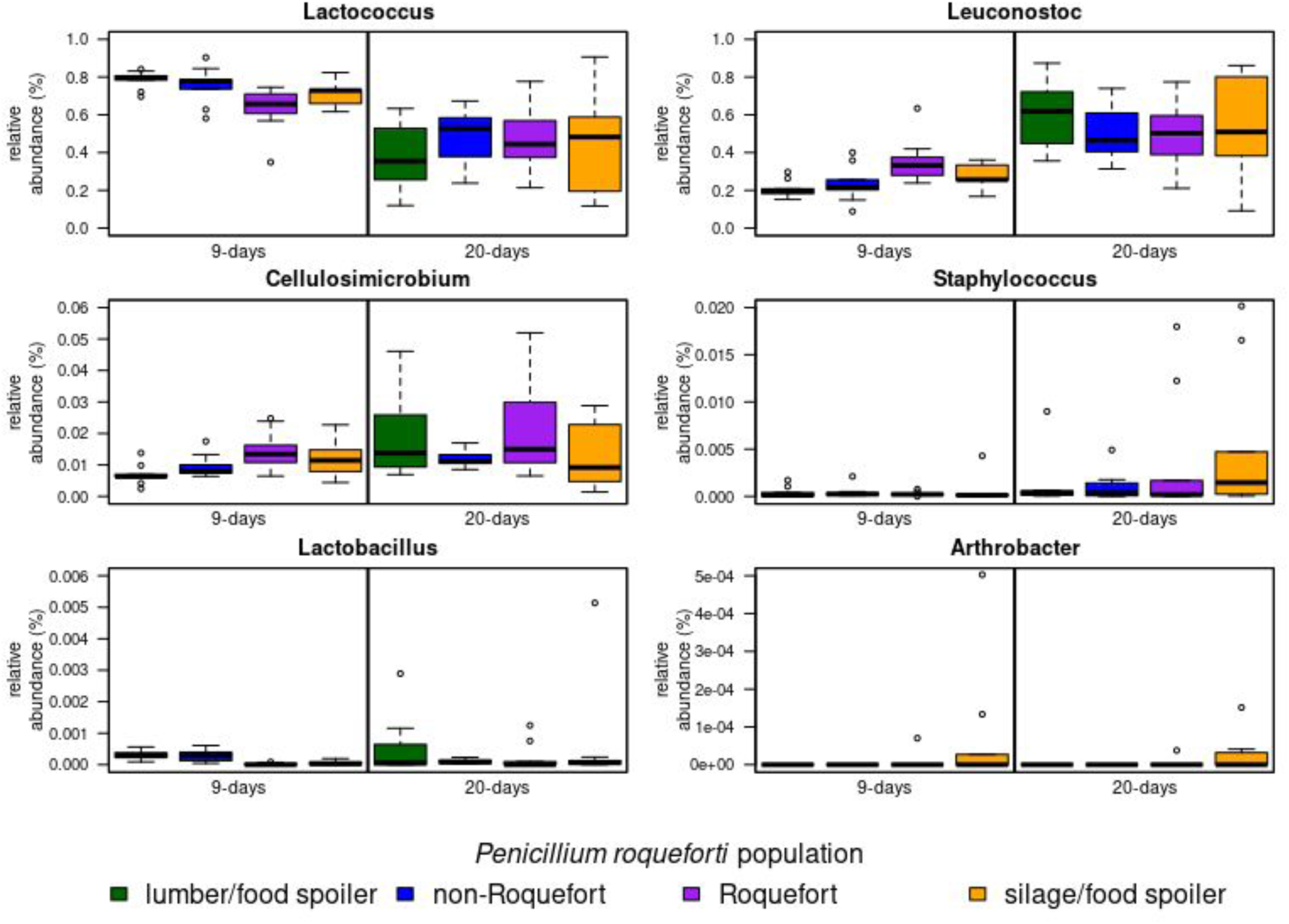
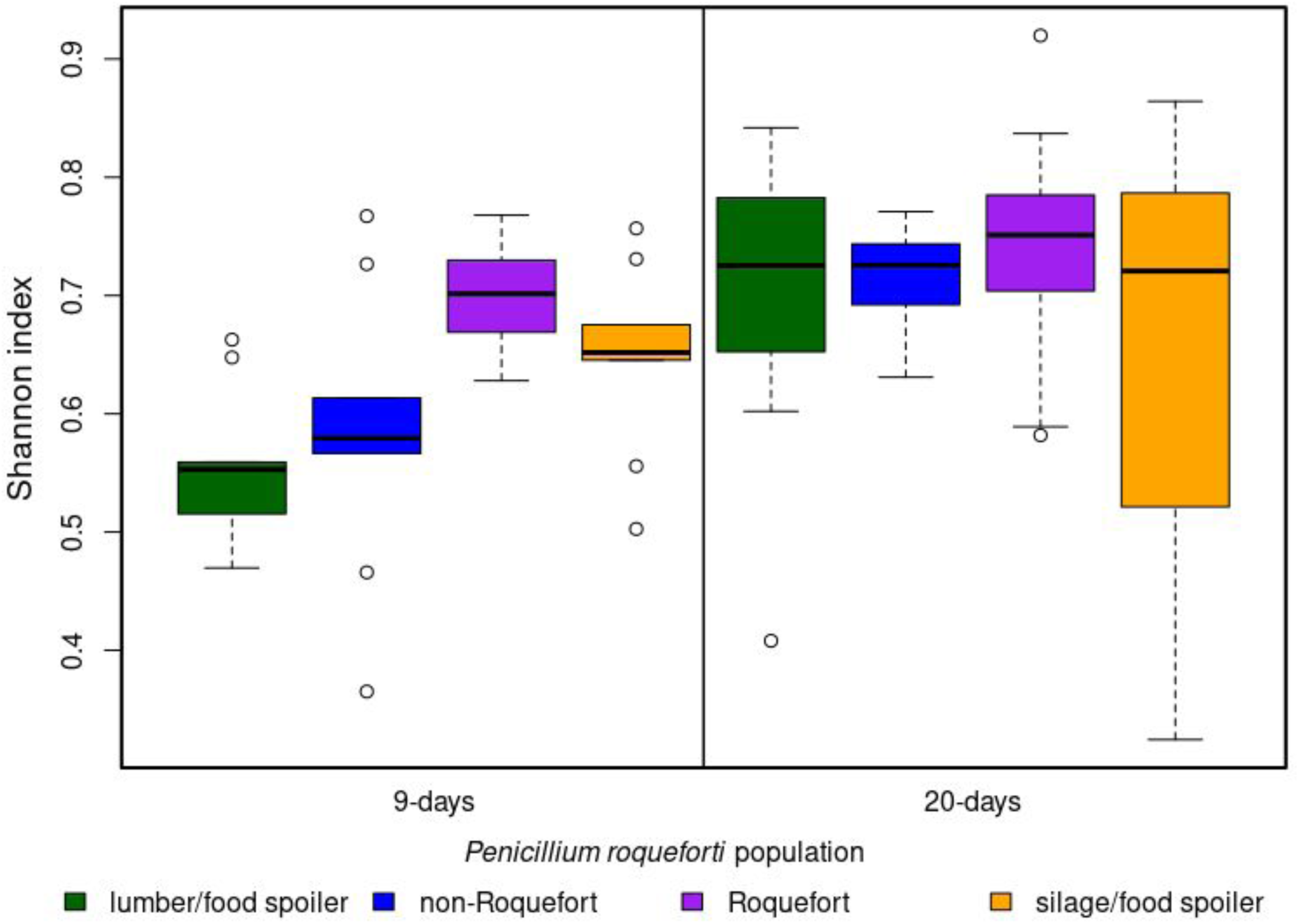
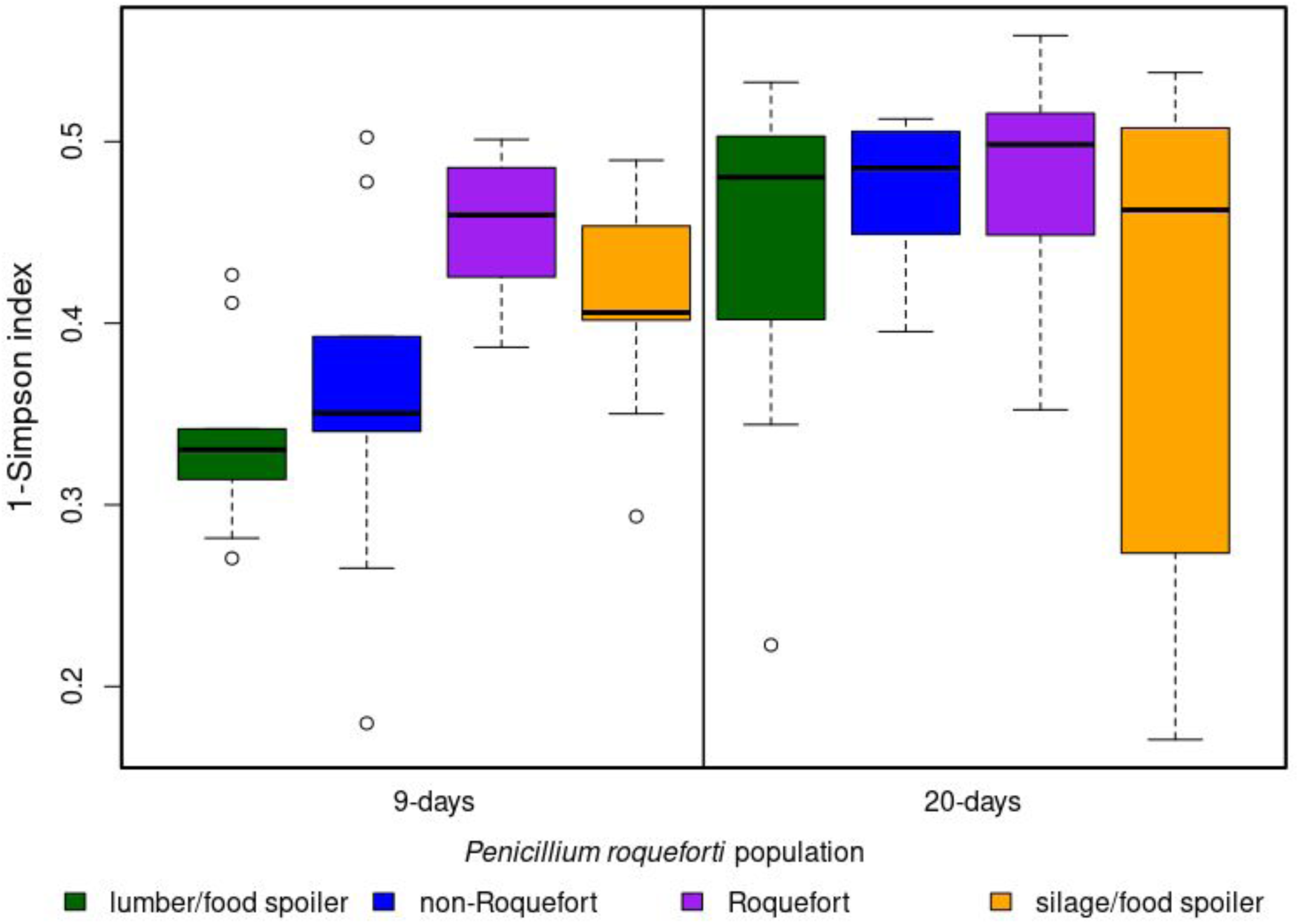
(A) Relative abundance of the six main bacterial operational taxonomic units in cheeses made with strains from the four *Penicillium roqueforti* populations (non-Roquefort cheese in blue, Roquefort cheese in purple, silage/food spoiler in orange and lumber/food spoiler in green) in cheeses at 9 (left) and 20 (right) days of maturation. (B & C) Mean bacterial genus diversity; A: Shannon index, B: Inverse of Simpson index (= 1 - Simpson index) for the operational taxonomic units detected by metabarcoding in 9-day cheeses (left) and 20-day cheeses (right) made with strains from the four *Penicillium roqueforti* populations (lumber/food spoiler in green, non-Roquefort cheese in blue, Roquefort cheese in purple and silage/food spoiler in orange).

### Higher proportion of blue area in cheeses produced with cheese *P. roqueforti* populations

The percentage blue area in cheese cavities was determined on freshly cut cheese slices (Figure 3B). The blue veins result from the formation of cavities in the cheese, and the growth and sporulation of *P. roqueforti* in these cavities. The percentage blue area was significantly higher in cheeses produced with cheese population strains than in those produced with non-cheese population strains (Figure 3A and 3B; Supplementary Table 1D). We also found a significant decrease in the percentage blue area from 20 to 180 days of maturation, for all populations except the Roquefort population, for which the percentage blue area remained relatively constant (Figure 3A; Supplementary Table 1D).

**Figure 3:**
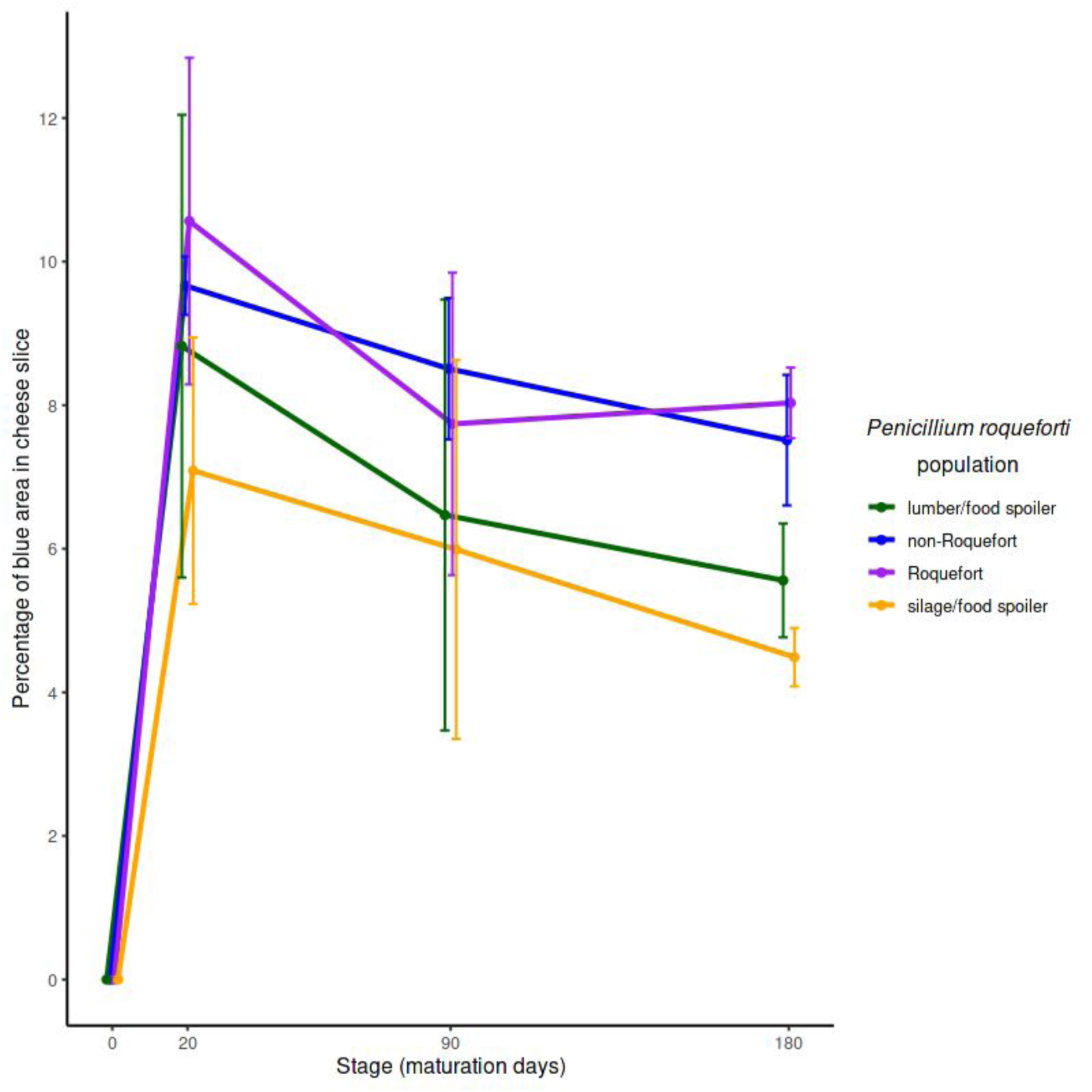
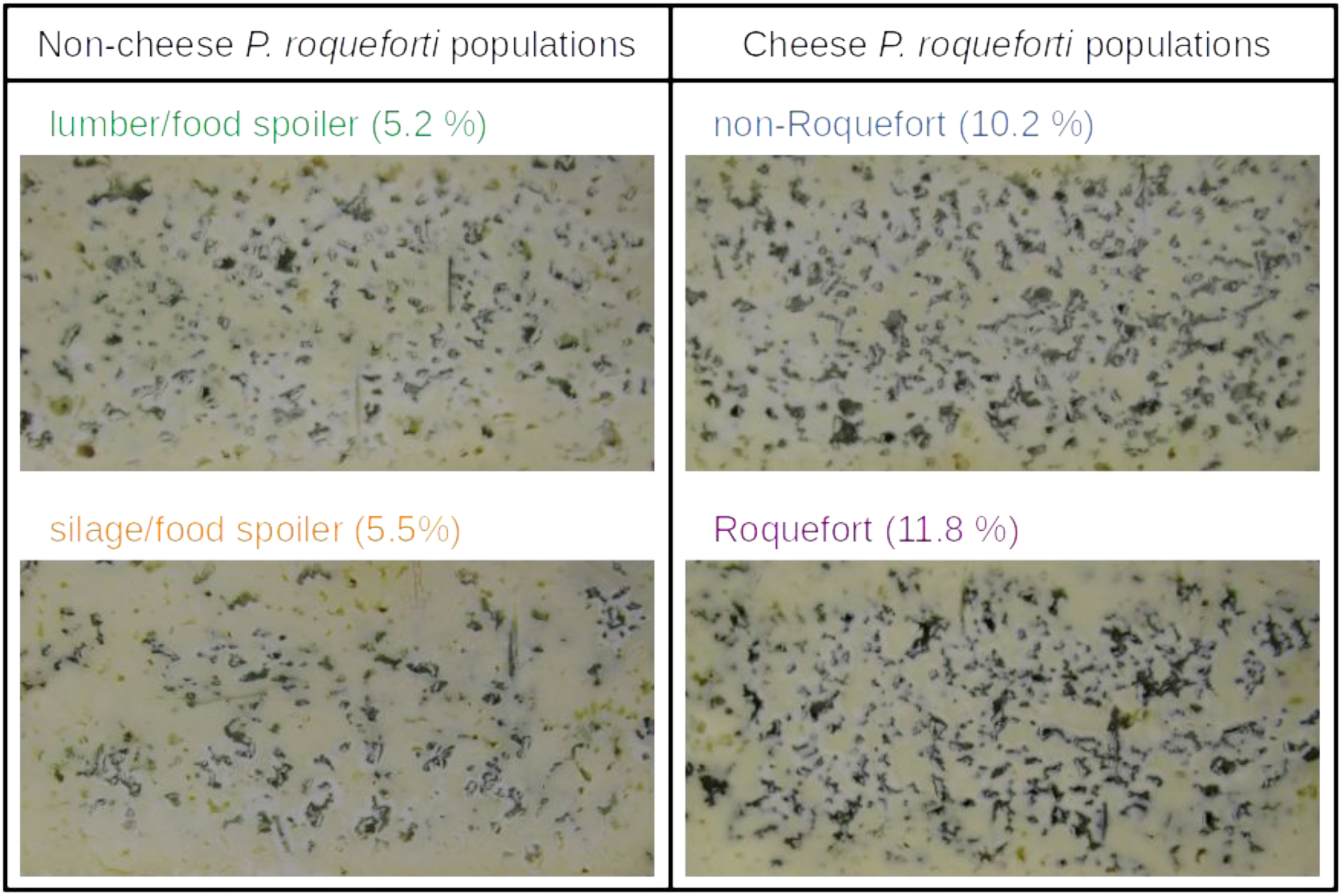
(A) Mean percentage blue area per cheese slice at 20, 90 and 180 days of maturation, for cheeses made with strains from the four *Penicillium roqueforti* populations (non-Roquefort cheese in blue, Roquefort cheese in purple, silage/food spoiler in orange and lumber/food spoiler in green). Error bars indicate 95% confidence intervals. (B) Illustration of the differences in mean percentage blue area per cheese slice at 180 days of maturation between the four *Penicillium roqueforti* populations (non-Roquefort cheese in blue, Roquefort cheese in purple, silage/food spoiler in orange and lumber/food spoiler in green). Contrast and brightness have been standardized and the edges cropped.

### More efficient proteolysis and lipolysis by the Roquefort *P. roqueforti* population

We investigated the proteolysis and lipolysis efficiencies of the four populations in cheeses after 90 days of maturation. Both targeted and non-targeted chromatographic analyses showed that proteolytic efficiency was highest in the Roquefort *P. roqueforti* population. We performed the targeted analysis with standards for the principal 23 amino acids (Supplementary Table 2A). We found that eight amino acids discriminated significantly between cheeses made with the different *P. roqueforti* populations (Supplementary Table 1E, mainly ornithine, leucine, and alanine), 15 discriminated between the cheese and non-cheese populations (mainly spermidine, isoleucine, methionine, glutamic acid, citrulline, serine and leucine) and 14 distinguished between the Roquefort and non-Roquefort populations (Supplementary Figure 3A, mainly valine, leucine, isoleucine, serine and threonine). The cheeses made with strains from cheese populations, and from the Roquefort population, in particular, had a higher total amino-acid concentration (Supplementary Tables 1E and 2B).

We also assessed proteolytic activity in a non-targeted analysis (fingerprint approach) on whole chromatograms (8,364 signals), which provided much more powerful discrimination between metabolites. Each metabolite generates a signal specific to its mass-to-charge (m/z) ratio at a given retention time. We obtained the largest number of aqueous signals, indicating the most efficient proteolysis, for cheeses inoculated with strains from the Roquefort population, followed by the lumber and non-Roquefort cheese populations, which were not significantly different from each other, and proteolysis was least efficient for the silage population (Figure 4; Supplementary Table 1F).

**Figure 4:**
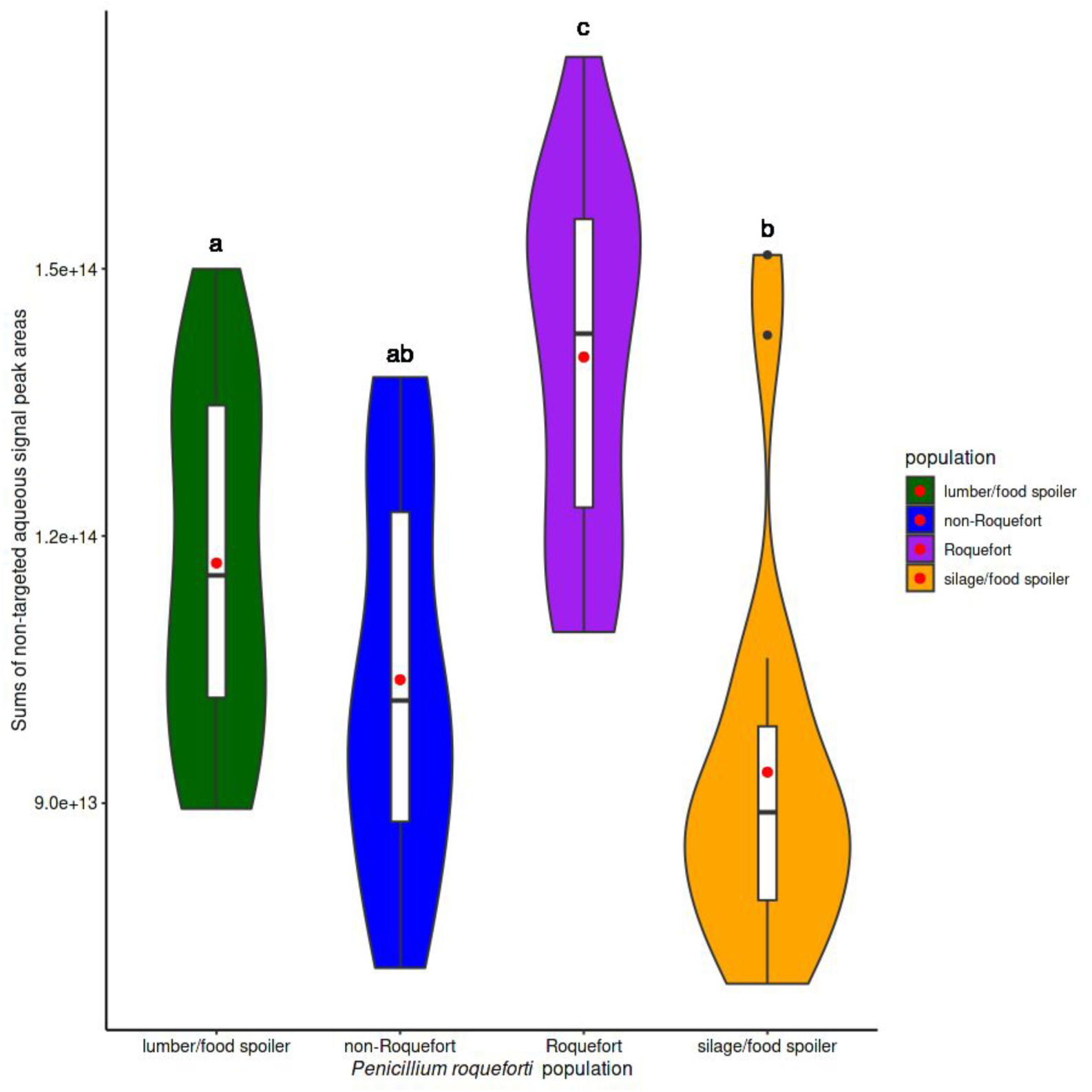
Violin plot depicting the distribution of the sums of 3,864 non-targeted aqueous signal peak areas, weighted by their mass-to-charge ratios (“m/z”), obtained in positive ionization mode for 90-day cheeses made with strains from the four *Penicillium roqueforti* populations (lumber/food spoiler in green, non-Roquefort cheese in blue, Roquefort cheese in purple and silage/food spoiler in orange). Boxplots within violin plots represent the median (center line), the 25th and 75th percentiles (box bounds), the 5th and 95th percentiles (whiskers), the points being the outliers from these 95th and 5th percentiles. The red dots depict the mean values.

Lipolysis was also more efficient for the Roquefort population than for the other populations. We investigated whether the *P. roqueforti* population influenced the abundance of free fatty acids and residual glycerides, as a proxy for the efficiency of lipolysis, in 90-day cheeses, with targeted and non-targeted chromatographic analyses in the positive and negative ionization modes. We specifically targeted glycerides and free fatty acids. In the targeted analysis, we identified seven free fatty acids and 20 triglycerides, and found that three free fatty acids (stearic, oleic and linoleic acids) were significantly more concentrated in cheeses made with Roquefort strains than in those made with strains from non-Roquefort populations (Supplementary Table 1G). In the non-targeted analysis, we obtained 3,094 signals and observed a higher abundance for organic signals specific to free fatty acids, indicating the most efficient lipolysis, in cheeses made with strains from the Roquefort population, followed by the lumber and non-Roquefort cheese populations, which were very similar to each other, with lipolytic efficiency lowest for cheeses made with strains from the silage population (Figure 5; Supplementary Table 1H). For residual glycerides, we obtained 8,472 signals, with no significant difference between the populations (Supplementary Figure 4; Supplementary Table 1I).

**Figure 5:**
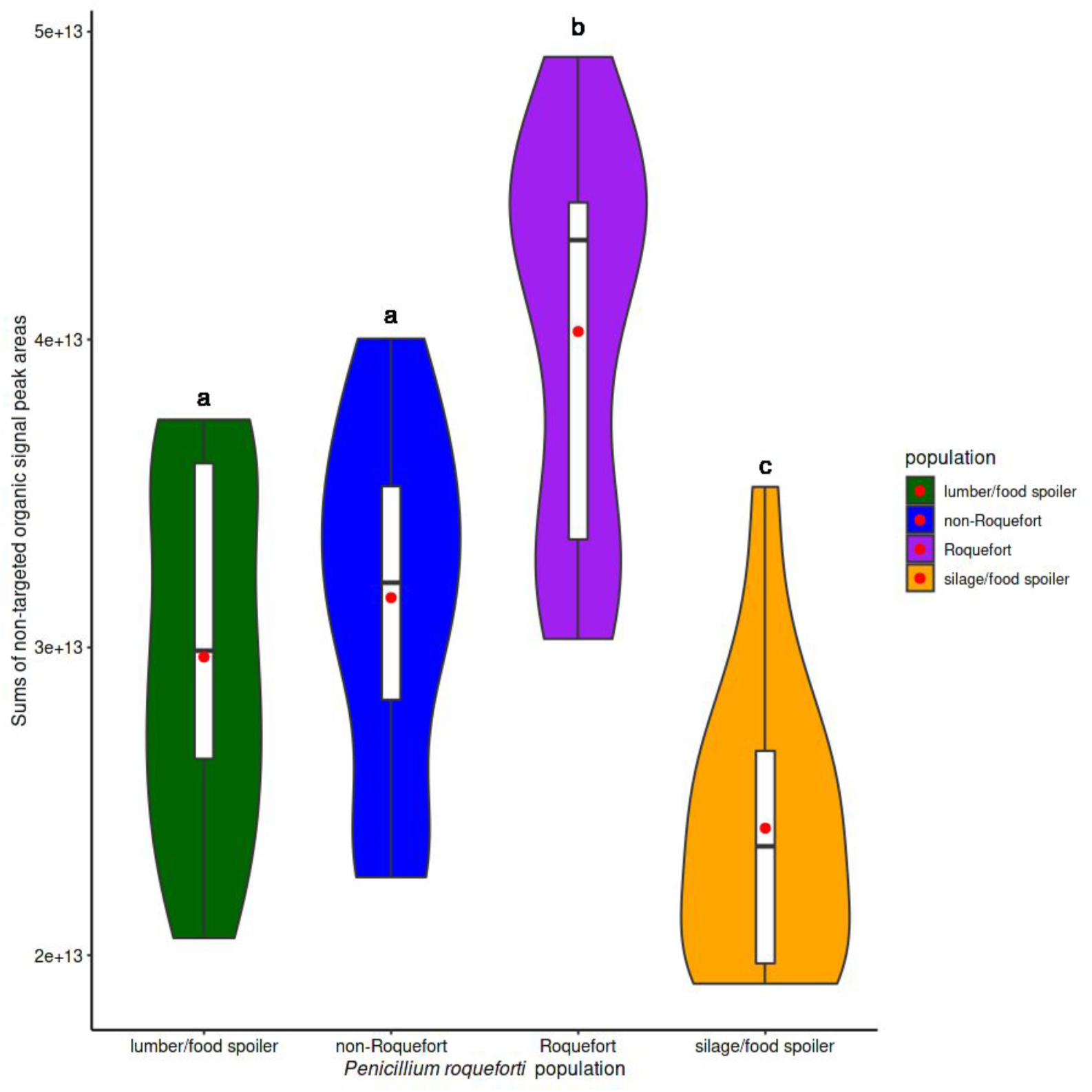
Violin plot depicting the distribution of the sums of 3,094 non-targeted organic signal peak areas, weighted by their mass-to-charge ratios (“m/z”), obtained in negative ionization mode for 90-day cheeses made with strains from the four *Penicillium roqueforti* populations (lumber/food spoiler in green, non-Roquefort cheese in blue, Roquefort cheese in purple and silage/food spoiler in orange). Boxplots within violin plots represent the median (center line), the 25th and 75th percentiles (box bounds), the 5th and 95th percentiles (whiskers), and the red dots depict the mean values.

We performed the most relevant standard physicochemical measurements on raw milk and on the cheeses at 9, 20, 90 and 180 days of maturation. As expected, we observed a maturation stage effect for 11 of the 16 physicochemical parameters (Supplementary Table 1J). Non-protein nitrogenous content was significantly higher in cheeses inoculated with strains from cheese *P. roqueforti* populations than in cheeses inoculated with strains from the other populations, consistent with the greater efficiency of proteolysis associated with these strains (Supplementary Figure 5A). Cheese water activity differed significantly between the cheeses made with strains from the four *P. roqueforti* populations (Supplementary figure 5B): it was significantly lower for the Roquefort cheese population than for the non-Roquefort cheese and silage populations (see statistics in the Supplementary Table 1J).

### Strong influence of *P. roqueforti* population on volatile compound production

We investigated the effect of *P. roqueforti* population on cheese volatile compounds in 90-day cheeses. We focused on the GC-MS data for the 40 principal volatile compounds considered to be most important for the aromatic quality of blue cheeses (Rothe *et al*., 1982): 11 acids, 12 ketones, 10 esters, six alcohols and one aldehyde (Supplementary Table 3). We found that *P. roqueforti* population strongly influenced the relative abundance of the compounds from these aromatic families in the cheeses (Supplementary Table 1K; Figures 6 and 7, and see below). In fact, the odors of the cheeses differed considerably (pers. obs.): the cheeses made with strains from the cheese *P. roqueforti* populations smelled as good as typical ripened blue cheeses, whereas those made with strains from non-cheese *P. roqueforti* populations had unpleasant odors, similar to those of a wet mop (Supplementary Figure 6; personal observation).

**Figure 6:**
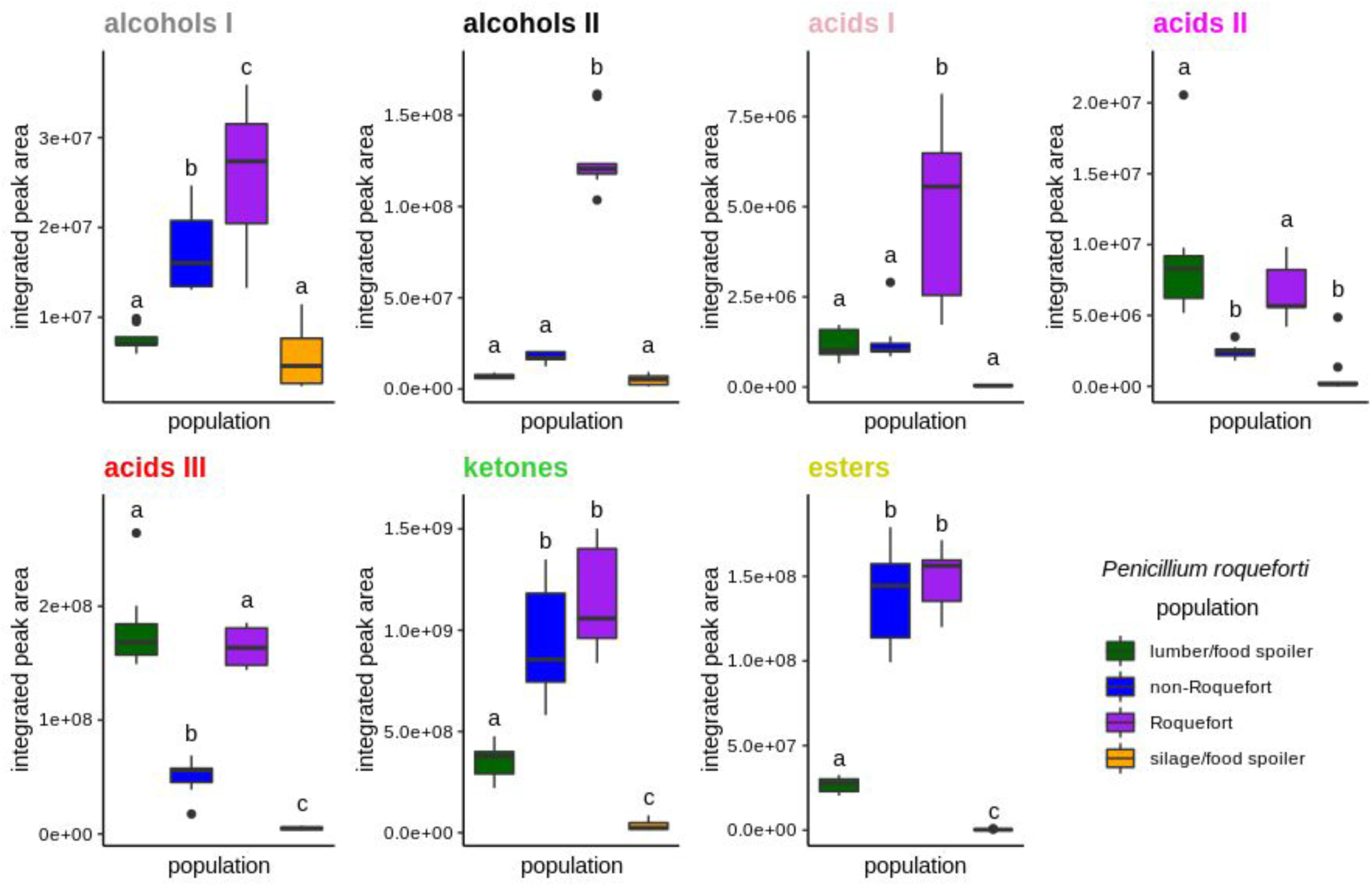
Volatile compound production (integrated peak areas from chromatograms in arbitrary units) in 90-day cheeses inoculated with strains from the four *Penicillium roqueforti* populations (non-Roquefort cheese in blue, Roquefort cheese in purple, silage/food spoiler in orange and lumber/food spoiler in green). The areas for each family of compounds are the sum of the integrated areas of the compounds belonging to the family concerned. Alcohols I and II are derived from proteolysis and lipolysis, respectively. Acids I, II and III are derived from proteolysis, glycolysis and lipolysis, respectively (Supplementary Table 3). The color of the titles indicates the affiliation of the compounds to their families, as in Figure S6.

**Figure 7:**
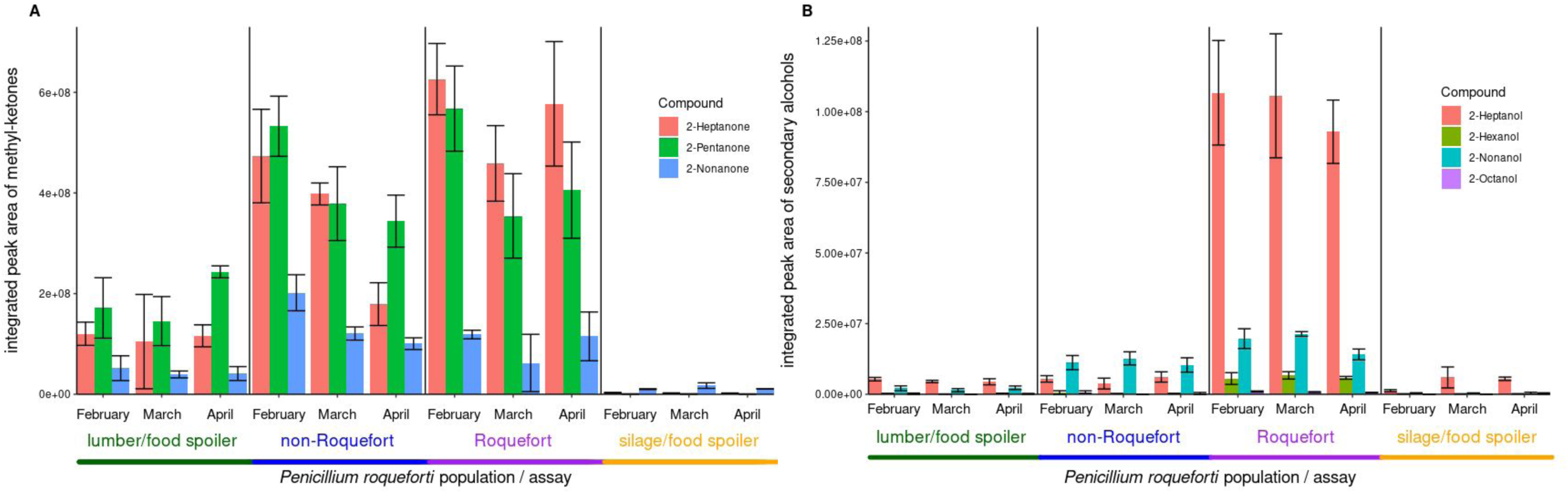
Integrated surface area (from chromatograms in arbitrary units) of the three most discriminant methyl ketones (A) and four secondary alcohols (B) for each assay (February, March, April) for the three strains of each *Penicillium roqueforti* population (lumber/food spoiler in green, non-Roquefort cheese in blue, Roquefort cheese in purple, silage/food spoiler in orange). Error bars represent standard deviations across cheese replicates.

The abundance of acids, methyl ketones and secondary alcohols resulting from proteolysis and lipolysis, and contributing to the typical flavor of blue cheese, was higher in cheeses produced with strains from cheese populations than in those produced with strains from non-cheese populations (Figure 6). These compounds were present at a particularly high abundance in cheeses made with strains from the Roquefort population. Four of the 40 compounds analyzed were proteolysis by-products (primary alcohols: 3-methyl-butanal, 3-methyl-butanol and isopropyl-alcohol, hereafter referred to as alcohols I, and 3-methyl-butanoic acid, hereafter referred to as acid I; Supplementary Table 3). The abundance of alcohols I was significantly higher in cheeses made with strains from cheese *P. roqueforti* populations than in those made with strains from non-cheese populations, and the highest values were obtained for the Roquefort population (mainly 3-methyl-butanol; Figure 6; Supplementary Table 1K). Acid I (i.e. 3-methyl-butanoic acid) was also present in higher abundance in cheeses made with strains from the Roquefort population than in other cheeses (Figure 6). Two acids, by-products of glycolysis (acetic and propionic acid, hereafter referred to as acids II), were present at higher abundance in cheeses made with strains from the Roquefort and lumber/food spoiler *P. roqueforti* populations than in other cheeses (Figure 6; Supplementary Tables 1K and 3). The other 35 aromatic compounds (i.e. acids from beta-oxidation, hereafter referred to as acids III, ketones, secondary alcohols hereafter referred to as alcohols II, and esters) were almost all direct or indirect by-products of lipolysis (Supplementary Table 3). The abundance of acids III was higher in cheeses made with strains from the Roquefort and lumber/food spoiler populations than in cheeses made with strains from the non-Roquefort cheese population (mainly butanoic, pentanoic, hexanoic and octanoic acid; Figure 6; Supplementary Table 1K). The levels of these compounds were lowest in cheeses made with strains from the silage population. Esters and methyl ketones (especially 2-pentanone and 2-heptanone) were present at higher abundance in cheeses made with strains from cheese *P. roqueforti* populations (Supplementary Table 1K). Cheeses made with strains from the Roquefort population contained the highest abundance of methyl ketones, and these compounds were barely detectable in cheeses made from silage population strains (mainly 2-heptanone, 2-pentanone and 2-nonanone; Figure 7A). The abundance of alcohols II, particularly 2-heptanol, was also much higher in cheeses made with Roquefort population strains than in other cheeses (Supplementary Table 1K; Figure 7B).

## Discussion

### Cheese *P. roqueforti* populations have been selected to produce better blue cheeses

Measurements of multiple features of blue cheeses made under conditions resembling those typically used in commercial Roquefort production revealed a strong influence of the differentiated *P. roqueforti* populations on several aspects of cheese quality. The cheese populations appeared the best adapted to cheesemaking, in terms of both the appearance and volatile compound content of the resulting cheese. The differences between the four *P. roqueforti* populations and the more appealing cheeses produced with strains from the cheese populations suggest that humans have exerted selection for the production of better cheeses, and this corresponds to domestication. Indeed, we found that cheese *P. roqueforti* strains produced a higher percentage blue area on cheese slices, an important visual aspect of blue cheeses. We also found that proteolysis and lipolysis were more efficient in cheeses made with Roquefort population strains than in cheeses made with strains from the other *P. roqueforti* populations, resulting in a higher abundance of desirable volatile compounds, including alcohols and associated acids. Cheese water activity was lower in cheeses made with strains from the Roquefort population, probably due to more efficient proteolysis (Ardö *et al*., 2017). We found no significant difference in the identities and abundances of microorganisms between the cheeses made with strains from the four *P. roqueforti* populations. Some minor differences in bacterial alpha diversity were observed, however, and the differences in all other measurements than diversity between cheeses probably reflected a direct effect of the specific features of the *P. roqueforti* population, although minor indirect effects involving the induction of more diverse bacterial communities by cheese *P. roqueforti* strains may also have occurred. Overall, our findings strongly support the view that cheese *P. roqueforti* populations have been selected by humans for better appearance and aroma. This selection may have involved the choice of the most beneficial strains for making good cheeses from standing variation, and/or the selection of *de novo* genetic changes. Previous studies found footprints of genomic changes in cheese populations in the form of beneficial horizontal gene transfers and positive selection (Dumas *et al*., 2020; Ropars *et al*., 2015).

Previous studies reported differences between *P. roqueforti* populations, in terms of growth, lipolysis and proteolysis, but on synthetic media (Dumas *et al*., 2020; Ropars *et al*., 2015). Here, using experimental cheeses made in commercial cheese production conditions, we reveal important features specific to cheese *P. roqueforti* populations, and to the Roquefort and non-Roquefort cheese populations. These findings are important in the context of domestication, for understanding rapid adaptation and diversification, and future studies based on quantitative trait mapping may be able to identify further genomic changes responsible for the specific features of the populations, according to the contrasting phenotypes revealed here. Progenies can indeed be obtained from crosses between strains from different populations of *P. roqueforti* (Ropars *et al*., 2015), and this could facilitate strain improvement through recombination between the different populations. Our results are, therefore, also important for improving blue cheese production.

### The four *P. roqueforti* populations induce similar microbiotas, but water availability is lower with cheese population strains, restricting the occurrence of spoiler microorganisms

Based on microbiological counts, we found no significant differences in abundance for any of the species monitored between cheeses made with strains from the four populations of *P. roqueforti*. In particular, we found no significant difference in the abundance of molds on Petri dishes. However, microbiological counts are known to provide poor estimates of fungal biomass, especially for mycelium growth (Schnurer, 1993).

The abundance and identity of the microorganisms studied were globally similar to those in four commercial Roquefort cheeses (personal information from C. Callon; Devoyod *et al*., 1968) and closely related blue cheeses (Diezhandino *et al*., 2015). The metabarcoding approach suggested that the different *P. roqueforti* populations induced bacterial communities of different levels of diversity. The cheese populations, and the Roquefort population in particular, were associated with the highest level of alpha diversity. The differences in cheese bacterial diversity, although minor, suggest that the differences between cheeses made with strains from the four *P. roqueforti* populations may be due not only to a direct effect of *P. roqueforti* population, but also to an indirect effect mediated by the induction of bacterial communities of different diversities. The large predominance of bacterial starters made it impossible to collect sufficient data for an assessment of the differences in relative abundance between subdominant bacterial species on the basis of metabarcoding. We also found a significant difference in water activity between cheeses made with strains from different *P. roqueforti* populations, the lowest value obtained being that for the Roquefort population. This may also reflect human selection, as low water activity restricts the occurrence of spoiler microorganisms. This characteristic is, therefore, subject to tight control in Roquefort cheeses for sale, particularly those for export.

### Cheese *P. roqueforti* populations produce bluer cheeses

We found significantly higher percentage blue areas in cheese slices from cheeses made with cheese *P. roqueforti* strains than in those made from non-cheese strains, potentially reflecting greater *P. roqueforti* growth in cheese and/or a higher sporulation efficiency in cavities. The percentage blue area in cheese slices also depends on the formation of cavities in the cheese, as *P. roqueforti* can only sporulate in cavities in which oxygen is available. The cavities are mostly generated by the gas-producing bacterium *Leuconostoc mesenteroides*, the abundance of which did not differ between the cheeses made with strains from different *P. roqueforti* populations, suggesting a direct effect of *P. roqueforti* populations on the blueness of cheese slices. The significantly higher percentage blue area in slices of cheeses made with cheese *P. roqueforti* strains than in those made with non-cheese strains therefore probably reflects better cheese and cavity colonization and sporulation, probably due to selection on the basis of appearance. The percentage blue area decreased by the end of maturation, perhaps due to the death of the fungus. Only cheeses made with Roquefort strains retained a high percentage blue area at 90 days of maturation, again potentially reflecting selection in pre-industrial times, when Roquefort cheeses had to be stored for several months at cave temperature before sale. The minimum maturation time for Roquefort PDO remains 90 days, which is longer than for other blue cheeses. These findings contrast with a previous study showing that a non-Roquefort population colonized the cavities of model cheeses better than other populations (Dumas *et al*., 2020); this discrepancy may reflect differences between studies in terms of the measurements used (total percentage blue area versus percentage blue area within cavities), the type of milk (ewe versus goat) or the mode of cheesemaking (rudimentary models versus commercial-like cheeses). Our findings are consistent with the presence of horizontally transferred genes in cheese populations with predicted functions in fungal development, including sporulation and hyphal growth (Dumas *et al*., 2020).

### Proteolysis and lipolysis are more efficient in the Roquefort *P. roqueforti* population

Based on chemical analyses and powerful chromatographic discrimination methods, we showed that the abundance of amino acids and small peptides (i.e., residual products of proteolysis) was highest in cheeses made with Roquefort *P. roqueforti* strains. Thus, these strains had the highest capacity for proteolysis, which is an important process in cheesemaking. Indeed, proteolysis contributes to the development of cheese texture, flavors and aromas (Andersen *et al*., 2010; Ardö, 2006, 2017; McSweeney, 1997; Roudot-Algaron, 1996). Previous measurements of proteolytic activity in synthetic media detected significant differences between *P. roqueforti* populations, but not between the two cheese populations (Dumas *et al*., 2020). We show here that experimental cheeses made with strains from the Roquefort population have a higher content of residual products of proteolysis, a sign of more advanced ripening.

We also found that lipolysis was more efficient in the cheeses made with strains from the Roquefort *P. roqueforti* population. By contrast, previous studies in synthetic media found that lipolysis was most efficient in the non-Roquefort population (Dumas *et al*., 2020). The discrepancy between these studies demonstrates the need for measurements in real cheeses for the reliable assessment of metabolic activities. Lipolytic activity is known to affect cheese texture and the production of volatile compounds affecting pungency (Alonso *et al*., 1987; González De Llano *et al*., 1990, 1992; Martín *et al*., 2016; Thierry *et al*., 2017; Woo *et al*., 1984). The more efficient proteolysis and lipolysis in the Roquefort *P. roqueforti* population should have a strong impact on cheese texture and flavor. It therefore probably results from selection to obtain better cheeses, i.e. from a domestication process, as previously reported for other fungi (Almeida *et al*., 2014; Baker *et al*., 2015; Gallone *et al*., 2016; Gibbons *et al*., 2012; Gonçalves *et al*., 2016; Libkind *et al*., 2011; Sicard *et al*., 2011). Roquefort cheeses are widely considered to be the blue cheeses with the strongest aromas and flavors. The less efficient lipolysis and proteolysis in the non-Roquefort population may result from more recent selection for milder cheeses.

### Cheese *P. roqueforti* populations produce cheeses with more abundant and diverse volatile compounds

We found major differences between the cheeses made with strains from different *P. roqueforti* populations, in terms of the volatile compounds resulting from lipolysis and, to a lesser extent, also from proteolysis. Only four of the aromatic compounds detected in our cheeses (3-methyl-butanal, 3-methyl-butanol, isopropyl-alcohol and 3-methyl-butanoic acid) were by-products of casein proteolysis (McSweeney *et al*., 2000), and the concentrations of these molecules were significantly higher in cheeses made with Roquefort *P. roqueforti* strains, consistent with the higher proteolysis efficiency and amino-acid precursor (i.e. valine, leucine and isoleucine) concentrations of these strains. These compounds produce fruity (banana), cheesy and alcoholic notes, which were probably important selection criteria during the domestication of the Roquefort *P. roqueforti* population. For the products of metabolic pathways leading from amino acids to alcohols (Ehrlich pathway with aldehyde reduction) or acids (aldehyde oxidation; Ganesan *et al*., 2017), the higher concentration of alcohols than of acids observed for all populations is consistent with the general micro-aerobic conditions of blue cheese cavities.

Most of the aromatic compounds identified were direct or indirect by-products of lipolysis, consistent with the known key role of lipolysis in the generation of typical blue cheese aroma (Cerning *et al*., 1987; Collins *et al*., 2003). The aromatic compounds resulting from lipolysis belonged to four chemical families (acids, methyl ketones, secondary alcohols and esters). Methyl ketones were the most diverse and abundant for cheese *P. roqueforti* populations, particularly for the Roquefort population, in which abundance was highest for 2-pentanone and 2-heptanone; 2-heptanone underlies the characteristic “blue cheese” sensory descriptor (Anderson *et al*., 1966; González De Llano *et al*., 1990, 1992; Moio *et al*., 2000). In *P. roqueforti,* methyl ketones with odd numbers of carbons are mostly produced by fatty-acid beta-oxidation, whereas those with even numbers of carbons may be produced by the beta-oxidation or autoxidation of fatty acids (Spinnler, 2011). These compounds are produced by the decarboxylation of hexanoic acid and octanoic acid, respectively, which were the most abundant acids found in our cheeses. This reaction is considered to be a form of detoxification, because methyl ketones are less toxic than acids (Kinderlerer, 1993; Spinnler, 2011). Interestingly, this pathway appeared to be more active in the cheese *P. roqueforti* populations, as methyl ketone levels were lower in cheeses made with lumber (four-fold difference) and silage (10-fold lower) strains than in cheeses made with cheese population strains. Methyl ketone concentrations were not directly associated with the concentrations of their precursors (acids), the highest concentrations being found in the lumber and Roquefort populations. The biosynthesis pathway producing methyl ketones must, therefore, be more efficient in cheese populations, particularly the non-Roquefort population. The cheese *P. roqueforti* populations were probably selected for their higher acid detoxification capacity, as this produces aromatic compounds with a very positive impact on flavor (Spinnler, 2011).

The concentrations of secondary alcohols (resulting from the reduction of methyl ketones) were also higher in cheeses produced by cheese *P. roqueforti* strains, particularly those of the Roquefort population, for which they were seven times higher than for the non-Roquefort cheese population and 20 times higher than for the silage/lumber populations; 2-heptanol was the major alcoholic compound produced. The reduction of 2-heptanone to 2-heptanol occurs specifically in anaerobic conditions and is much stronger in the Roquefort population; aerobic conditions were similar for all the populations. The Roquefort *P. roqueforti* population may also have been selected for this feature, as secondary alcohols provide “fruity notes”, which are associated with better aromatic quality (Spinnler, 2011). Methyl ketones may be reduced to alcohols by an alcohol dehydrogenase, as occurs when aldehyde is reduced to alcohol via the Ehrlich pathway. Alcohol dehydrogenase genes may thus have been targets of selection in the Roquefort *P. roqueforti* population, although they were not detected as evolving under positive selection in a previous study (Dumas *et al*., 2020).

We also found higher levels of esters in cheeses made with cheese *P. roqueforti* populations. Esters are produced principally by the esterification of ethanol with acids generated by beta-oxidation. *Leuconostoc* starters can produce ethanol, and ester synthesis has also been described as a detoxification mechanism (Mason *et al*., 2000). These results further indicate that cheese *P. roqueforti* populations, particularly the Roquefort population, have been selected for acid detoxification capacity, leading to a large variety of less toxic aromatic compounds with strong aromas and flavors.

Overall, the aromas of cheeses made with cheese *P. roqueforti* strains had more appealing aromas, and this was particularly true for cheeses made with Roquefort strains. These aroma properties probably reflect selection by humans. The cheeses made with silage and lumber populations had a mildly unpleasant smell, whereas those made with cheese strains smelled like typical blue cheeses, with cheeses made with Roquefort strains having the strongest smell. This may reflect previously reported horizontal gene transfers in cheese populations, involving genes with predicted functions in lipolysis or amino-acid catabolism, and the positive selection of genes involved in aroma production (Dumas *et al*., 2020). We compared *P. roqueforti* populations between cheeses made following commercial modes of production, which represents a major advance relative to previous studies based on experimental models or synthetic media (Dumas *et al*., 2020; Gillot *et al*., 2017). We used unpasteurized ewe’s milk, in accordance with the requirements for Roquefort PDO production, which also affects cheese aromas. In future studies, it would be interesting to determine whether the use of pasteurized or unpasteurized ewe’s milk or cow’s milk leads to similar specific features of the Roquefort versus non-Roquefort cheese *P. roqueforti* populations, as there may have been selection during domestication, leading to an adaptation of the Roquefort population for the catabolism of unpasteurized ewe’s milk.

### Conclusion

We show that the *P. roqueforti* population has a strong impact on cheese quality, appearance and aroma. The populations used for cheesemaking led to bluer cheeses, with better aromas, probably due to domestication involving the selection of multiple fungal traits by humans seeking to make the best possible cheeses. French cheese producers have been inoculating cheeses with *P. roqueforti* spores from moldy rye bread since the end of the 19^th^ century (Labbe *et al*., 2004, 2009; Vabre, 2015). This process made it possible for them to re-inoculate with the strains producing the best cheeses, thereby applying a strong selection pressure. The two cheese populations displayed a number of specific features, with the Roquefort population notably producing more intense and specific aromas and flavors. The selection of different fungal varieties for different usages has also been reported in the fermenting yeast *Saccharomyces cerevisiae* (Gallone *et al*., 2016; Legras *et al*., 2018). Previous studies on *P. roqueforti* detected recurrent changes in amino acids and horizontal gene transfers in cheese populations, both of which facilitated rapid adaptation (Dumas *et al*., 2020; Ropars *et al*., 2015). Our findings provide greater insight into *P. roqueforti* domestication and pave the way for strain improvement through the targeting of relevant traits. A protocol inducing sexual selection has been developed in *P. roqueforti* (Ropars *et al*., 2014), making it possible to perform crosses between strains from the two cheese populations, each of which harbors very little genetic diversity (Dumas *et al*., 2020), to generate variability and to identify strains with high levels of performance. The results of this study will facilitate the choice of the parental strains for crossing and of the most important phenotypes to be measured in the offspring. Parental strains with strongly contrasting phenotypes for the traits important for cheesemaking that we found to be differentiated between populations (such as volatile compound production, lipolysis and proteolysis) should be used, to maximize variability in the progeny. Other traits may also be worth investigating in the future, to understand the changes that have occurred during the domestication of the two cheese *P. roqueforti* populations, particularly as concerns toxin production in cheeses. Previous *in vitro* studies have shown a lack of production of the toxic mycophenolic acid by the non-Roquefort population due to a deletion in one of the genes of the synthesis pathway (Gillot *et al*., 2017). In the *P. camemberti* fungus, domestication has also led to a lack of production of a mycotoxin due to a deletion in a gene (Ropars *et al*., 2020b).

## Acknowledgments

We thank Béatrice DESSERRE, Céline DELBES and Cécile CALLON for advice and technical assistance in microbiology, Sébastien THEIL for technical support for metabarcoding analyses, Patricia LE THUAUT, Manon SURIN and Brigitte POLLET for technical support for metabolomic analyses, Sara PARISOT for milk delivery and quality, Pierre CONCHON for technical support for image analysis, LIAL-MC for the various reference measurements in physicochemistry, Christophe LACROIX and Alfonso DIE for the determination of short-chain fatty acids in fermentation supernatants.

This study was funded by the LIP SAS, ANRT (*Association Nationale Recherche Technologie*), the ERC Genomefun 309403 starting grant, the BLUE ERC Proof-of-Concept grant, the Fondation Louis D grant (French Academy of Sciences) and the ANR-19-CE20-0002-02 Fungadapt ANR grant.

## Figure legends for the Supplementary Material

Figure S1: Abundance of microorganisms (in log colony-forming units/g) of the eight types monitored at various stages of cheese maturation (i.e. unpasteurized milk, 9, 20, 90 and 180 days), for each of the four *Penicillium roqueforti* populations used to inoculate the cheeses (non-Roquefort cheese in blue, Roquefort cheese in purple, silage/food spoiler in orange and lumber/food spoiler in green). Error bars represent standard deviations across assays.

Figure S2: Illustration of image processing for estimation of the percentage blue area on cheese slices: (a) example of an unprocessed image of a cheese slice; (b) image after brightness and contrast standardization; (c) image after cropping; (d) corresponding image binarization with a grayscale of 102 on the red channel. White and black correspond to pixel classification: in white, the inner part of the cheese and empty cavities; in black, cavities filled with the fungus.

Figure S3: Differences in aminoacid content between cheeses according to the population-of-origin of the *Penicillium roqueforti* strains. A. Discrimination between 90-day cheeses made with cheese (blue) and non-cheese (green) *P. roqueforti* populations (left), or Roquefort cheese (purple) and non-Roquefort cheese (blue) *P. roqueforti* populations (right), based on the abundance of the 23 identified amino acids present, according to an orthogonal signal-corrected partial least squares (PLS) discriminant analysis. Vertical and horizontal axes represent PLS1 and PLS 2 scores and gray arrows represent the relative contribution of loadings of signals significantly discriminating the group considered in a *t*-test with jackknife resampling. B. Abundance of molecules from particular classes detected in cheeses: mean integrated peak area from chromatograms in arbitrary units (bars, left axis) and cumulative percentage (line with dots, right axis) of aqueous extracts across all 90-day cheeses.

Figure S4: Violin plot depicting the distribution of the sums of 8,472 non-targeted organic signal peak areas, weighted by their mass-to-charge ratios (“m/z”), obtained in positive ionization mode for 90-day cheeses made with strains from the four *Penicillium roqueforti* populations (lumber/food spoiler in green, non-Roquefort cheese in blue, Roquefort cheese in purple and silage/food spoiler in orange). Boxplots within violin plots represent the median (center line), the 25th and 75th percentiles (box bounds), the 5th and 95th percentiles (whiskers), and the red dots depict the mean values.

Figure S5: Non-protein nitrogen levels at 20, 90 and 180 days of maturation, and water activity at 90 and 180 days of maturation. Comparison of cheeses made with strains from different *Penicillium roqueforti* populations (non-Roquefort cheese in blue, Roquefort cheese in purple, silage/food spoiler in orange and lumber/food spoiler in green). Error bars indicate 95% confidence intervals.

Figure S6: Discrimination between 90-day cheeses inoculated with strains from the four *Penicillium roqueforti* populations (non-Roquefort cheese in blue, Roquefort cheese in purple, silage/food spoiler in yellow and lumber/food spoiler in green), based on the abundance of 41 volatile compounds in an orthogonal signal-corrected partial least squares (PLS) discriminant analysis. Vertical and horizontal axes represent the PLS1 and PLS2 variances, and arrows represent the relative contributions of compound odor loadings significantly discriminating the group considered (according to www.thegoodscentscompany.com) in a *t*-test with jackknife resampling. The odor colors indicate the families in Figure 6 to which the associated compounds belong.

