## Supplementary figures and images for "Strong effect of *Penicillium roqueforti* populations on volatile and metabolic compounds responsible for aromas, flavor and texture in blue cheeses"

### Figure S1

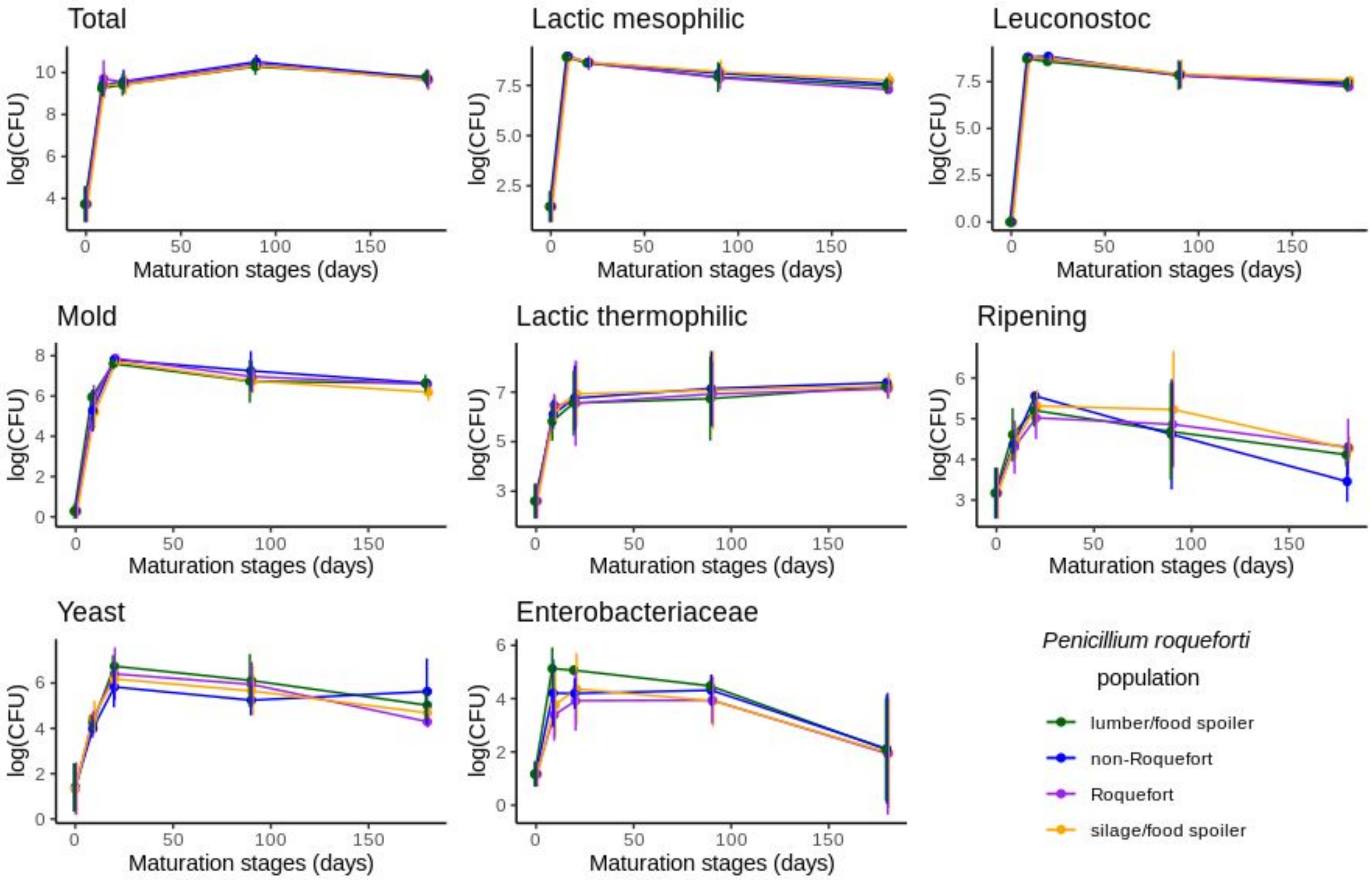

### Figure S2

Figure S2

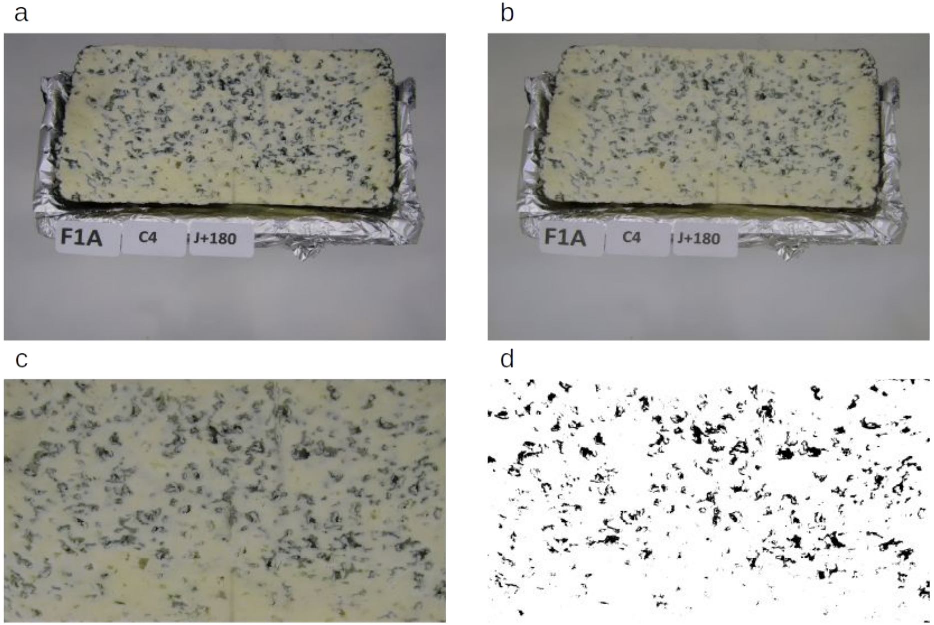

### Figure S3A

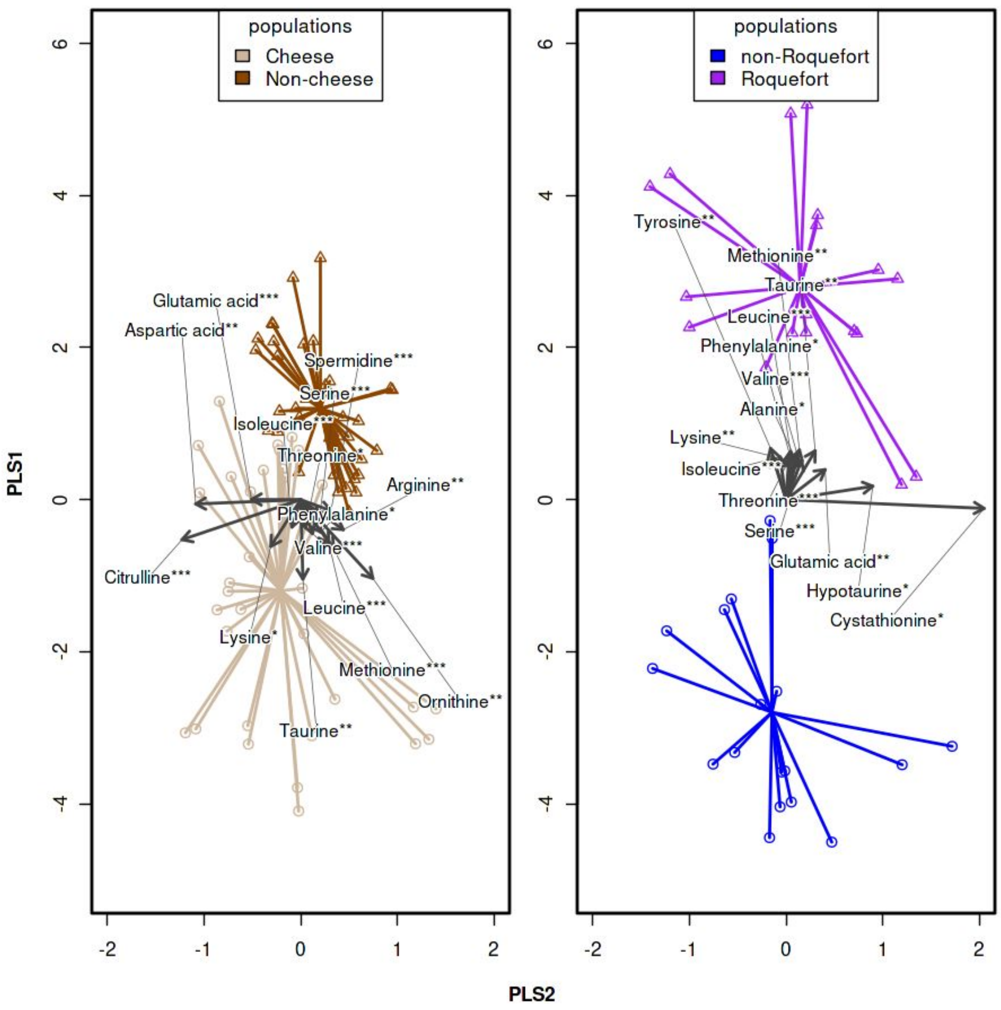

### Figure S3B

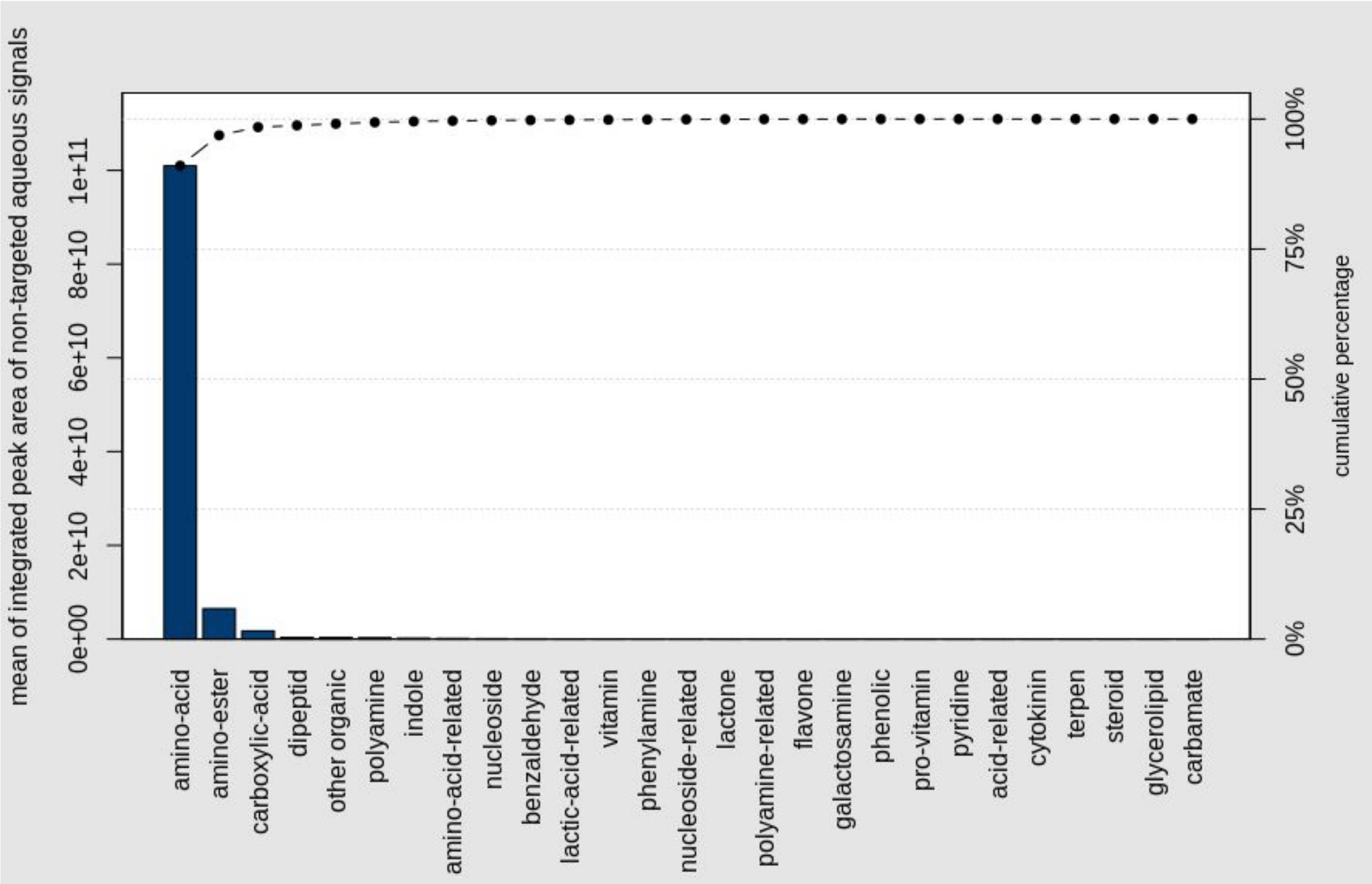

### Figure S4

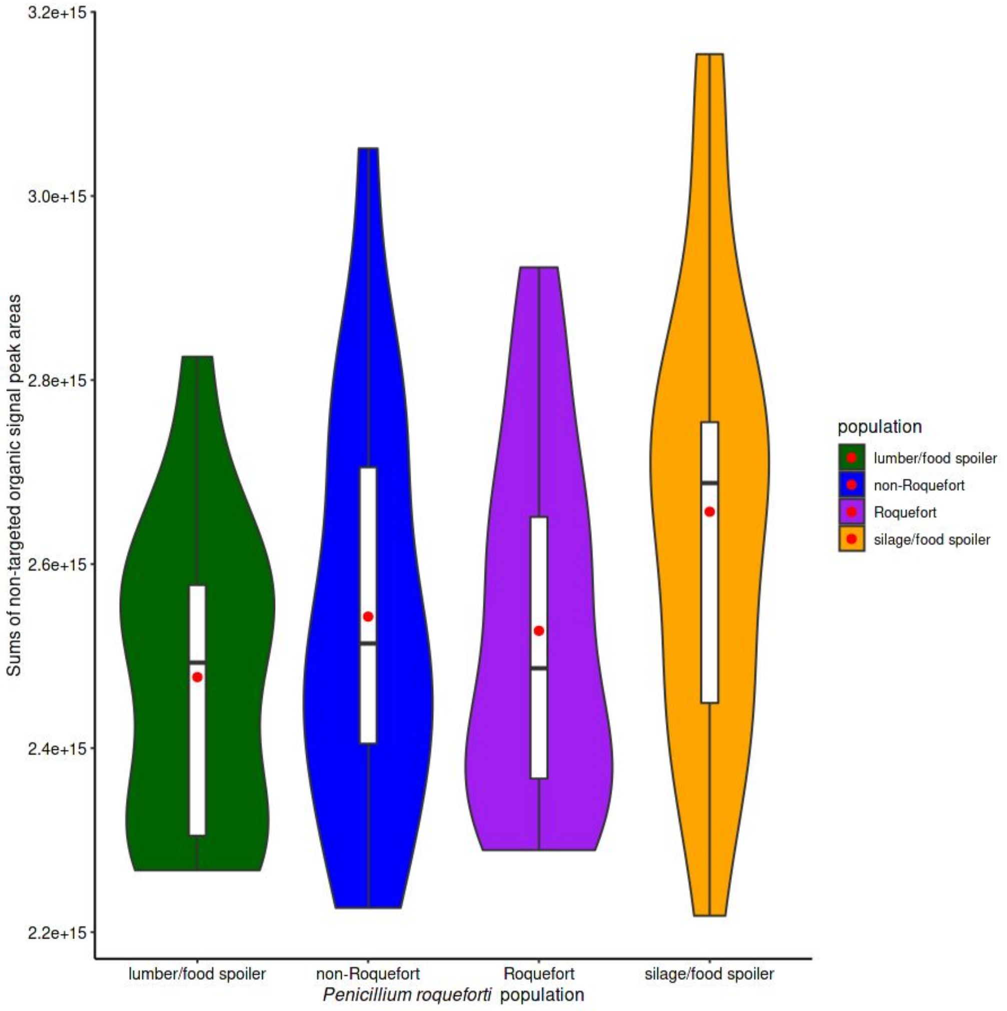

### Figure S5

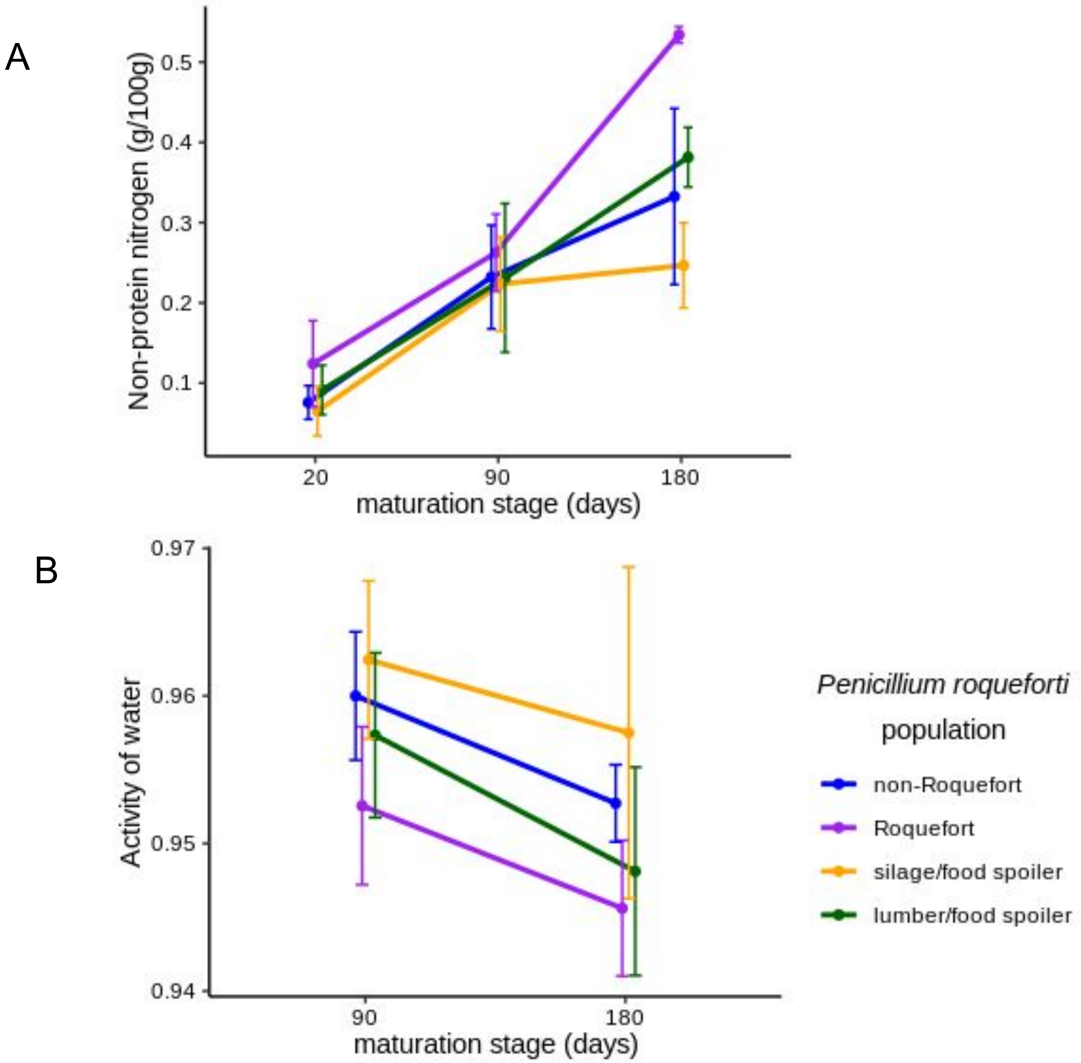

### Figure S6

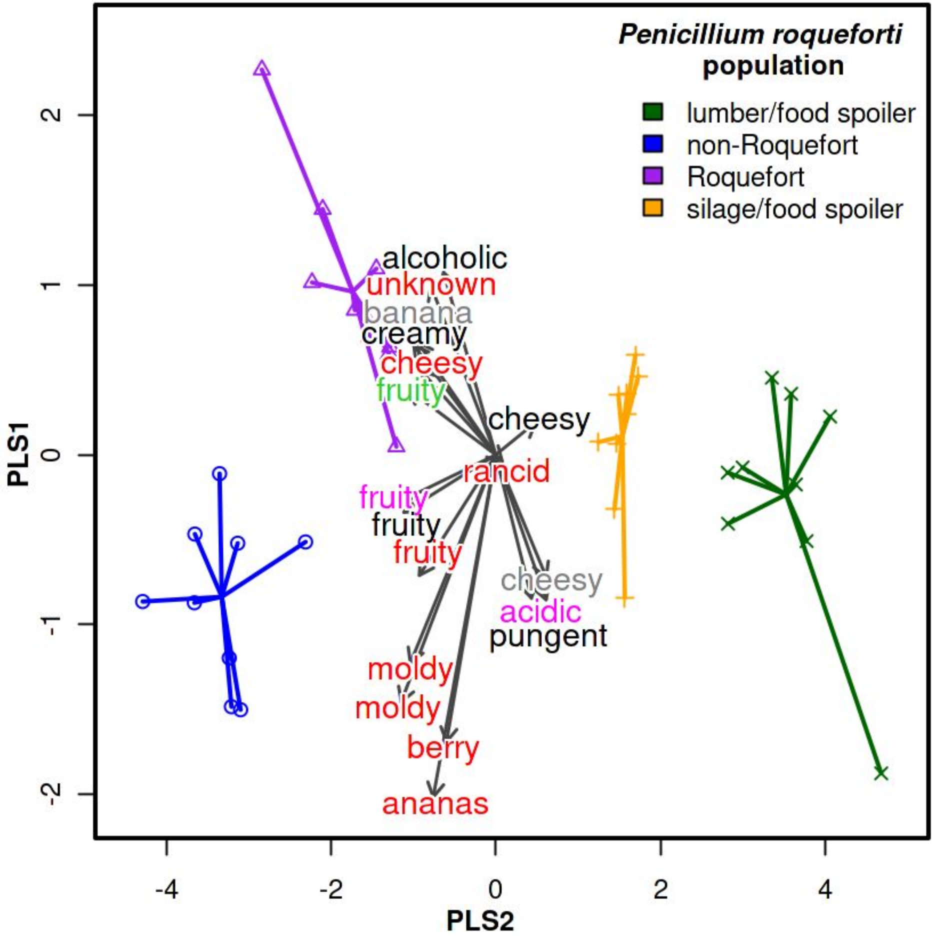
