## Supplementary Methods for "Strong effect of *Penicillium roqueforti* populations on volatile and metabolic compounds responsible for aromas, flavor and texture in blue cheeses"

**Materials and methods**

1. **Experimental design and fungal strains**

We made cheeses using three different strains of each of the four *P. roqueforti* populations (Figure 1), assigned to the populations in a previous study based on molecular markers (Dumas *et al.*, 2020). Because of production capacity limitation in the experimental facility, we could not make all cheeses at the same time. We therefore divided the production into three assays, each including one strain from each of the four populations (Figure 1A). For each strain within each assay, we made three production replicates, with two cheeses per strain in each replicate to have enough material to sample, amounting to a total of 72 cheeses (4 strains * 2 cheeses * 3 replicates * 3 assays). Because the assays were made sequentially from February to April, the effect of the seasonal change in milk composition was confounded with the strain effect within the population, hereafter referred to as the "assay effect". The three replicates within each assay were also done at different times, and thus with different batches of raw milk (Figure 1A). LCP strains (“Laboratoire de Cryptogamie de Paris”) are available from the Muséum d’Histoire Naturelle de Paris, France, and BRFM strains (“Biological Ressource Fungi Marseille'') are available from the Centre International de Ressources Microbiennes de Marseille, France.

2. **Cheesemaking**

The cheesemaking protocol corresponded to the typical mode of production used by the main Roquefort producers and complied with the Roquefort protected designation of origin (PDO) specifications, except that the ripening process occurred in artificial cellars in Aurillac INRA facilities and with strains of different *P. roqueforti* populations. For each cheesemaking production assay, 150 L of raw Lacaune sheep milk from the La Fage experimental farm (43°55’6.877’’N, 3°5’40.806’’E, Menet, France) was collected and stored at 4°C for one night. Technological parameters (temperature, pH, rennet clotting time and dry matter content of milk, curd and cheese) were monitored during cheesemaking production. In each production assay, performed in Aurillac INRA facilities, one vat was used per strain, each vat being used for making two cheeses and cleaned between each cheesemaking. The manufacturing schedules for the different vats were staggered by 10 minutes intervals. In a room at 26°C / 98% humidity, 30 to 35 L of milk were slowly mixed in each vat and heated up to 32.5°C. Then, we added 20 mg.L^-1^ of mesophilic starter culture with *Leuconostoc* spp., *Lactococcus lactis* subsp. *cremoris*, *L. lactis* subsp. *lactis*, and *L. lactis* subsp. *lactis* biovar *diacetylactis* (FD-DVS CHN-11, CHR HANSEN, Saint-Germain-lès-Arpajon, France), 25 mg.L^-1^ of gas-producing *Leuconostoc mesenteroïdes* subsp. *mesenteroïdes* (CHOOZIT LM 57 LYO 20 DCU, DANISCO, Paris, France) and ca. 3.6e6 UFC.L^-1^ of *P. roqueforti* spores (in 20% salt water, SAS LIP, Aurillac, France). After five minutes of homogenization, we added 0.25 ml.L^-1^ of active Chymosin at 520 mg.L^-1^ (Laboratoires Humeau, La Chapelle-sur-Erdre, France). After three minutes of homogenization, the milk was left curing for 110 min. The curd was cut into 1.5 cm^3^ cubes, left to rest for five minutes, and brewed for 50 minutes. After draining, the curd was put in drilled cylinders of 20x10 cm at 26°C / 85% humidity, and cheeses were returned five times at 10, 20, 60, 120 and 240 minutes after moulding. The first day after cheesemaking, the room was set at 23°C, cheeses were turned twice, and then once a day until the 20th day. On the second day, cheeses were unmoulded. On the third day, cheeses were rubbed with sterilised coarse salt and then transferred into ripening cellars at 11°C / 95% humidity. On the fifth day, cheeses were salted again the same way. On the seventh day, cheeses were pricked with 37 picks distributed over the entire surface to form holes in cheeses where oxygen can allow *P. roqueforti* growth and sporulation within the cheeses. The ripening period ended on the twentieth day, when cheeses were wrapped in 45x45 cm sterile aluminum foil and then aged at -2°C until 180^th^ day.

Cheese samples were collected on days 0, 9, 20, 90 and 180, referred hereafter as “stages”. Day 0 corresponded to raw milk for microbiology measures and sowed curd for metabolic and volatile compound analyses. Days 9 and 20 corresponded to the half and end of the ripening period, respectively. Days 90 and 180 (i.e. 3 and 6 months) corresponded to the minimum maturing time for Roquefort PDO and to a typical sale stage of ageing, respectively.

3. **Microbiological analyses**
   a. **Microbiological counts**

We estimated the concentration of different microorganisms floras on the initial crude milk and on every sampled cheese maturation stage to test if the *P. roqueforti* population had an influence on cheese microbiota. Microbiological counts were performed on raw milk while cheeses were frozen at -20°C for further analysis. A total of 10 g of cheese, including rind and core, were added to 90 mL phosphate buffer pH 7.5 (Gomri, 1946) and homogenized for four minutes in a stomacher (AES, Lutzelbourg, France). These milk and cheese suspensions were diluted with sterile Ringer’s solution and plated on specific media. Total aerobic mesophilic bacteria number was counted after three days at 30°C on plate count agar (PCA ; Nelson, 1940). Mesophilic lactic acid bacteria were enumerated on Man Rogosa Sharpe agar (MRS) incubated at 30°C for 3 days (De Man *et al.*, 1960) and thermophilic lactic acid bacteria on M17 agar (Terzaghi, 1975) incubated à 42°C for two days. Dextran-positive *Leuconostoc* spp. were enumerated on Mayeux, Sandine and Elliker medium (MSE; Mayeux *et al.*, 1962) after 2 days at 30°C. Molds and yeasts were enumerated on oxytetracycline gelose agar medium (OGA; Mossel *et al.*, 1970) after 3 days at 25°C. Gram-positive catalase-positive bacteria were enumerated on cheese-ripening bacterial medium (CRBM; Denis *et al.*, 2001) incubated 10 days at 25°C. Enterobacteria were enumerated on violet red bile glucose (VRBG; Mossel *et al.*, 1978) incubated in anaerobic conditions during 24h. All media were purchased from Biokar Diagnostics (Biokar, Beauvais, France).

b. **Metabarcoding analysis**

In order to further test whether the *P. roqueforti* population had an effect on the identity and relative abundance of other microorganisms in cheeses, we performed a metabarcoding analysis on our experimental cheeses at 9 and 20 days (ripening period), by extracting DNA, amplifying and sequencing a DNA fragment (16S) typically used for bacteria molecular identification. DNA extraction was performed on cheeses kept at -20°C following Duval *et al.*, 2016 with some modifications. A total of 15 g of cheese, including rind and core, were added to 4 mL of sterile water and homogenized for 2 minutes in a stomacher (Interscience, St-Nom la Bretèche, France). A total of 250 mg of this homogeneisate was put into a grinding tube (Sarstedt, Nümbrecht, Germany) with 250 μL of 4M guanidium thiocyanate/0.1M Tris-HCl (pH 7.8), 40 μL of 10% N-laurylsarcosine in deionized water (Milli-Q Reagent Grade Water System, Millipore Corporation, Billerica, Massachusetts, USA) and 200 mg of silica-zirconium (50/50 of 0.1 mm and 0.5 mm diameter, BioSpec, Bartlesville, USA). Suspensions were shaken 20 seconds at 6,500 m.s^-1^ in a Precellys Evolution (Bertin Instruments, Montigny-Le-Bretonneux, France). Then, 75 μL of a mix with 3 mg lysozyme and 20 μL lyticase (5000 U.mL^-1^) in 55 μL of Tris-EDTA-sucrose were added and vortexed. Suspensions were heated at 32°C for 30 min and vortexed for 15 min. Next, 40 μL of 14 mg.mL^-1^ proteinase K (VWR, Fontenay-sous-Bois, France) and 100 μL of 20% sodium dodecyl sulfate were added and vortexed. Suspensions were heated at 55°C for 30 min and vortexed for 15 min. To 250 μL of this mixture were added 200 μL of sodium phosphate buffer, 200μL of sodium acetate-EDTA solution and 500μL of phenol:chloroform:isoamyl alcohol at 25:24:1 (pH 8; VWR, Fontenay-sous-Bois, France). Sodium phosphate buffer was made with 1.33% of Na_2_HPO_4_.12H_2_O and 0.07% of NaH_2_PO_4_.H_2_O at 0.2M. Sodium acetate-EDTA solution was made with 0.68% of sodium acetate and 0.37% EDTA. Suspensions were shaken for 45 seconds at 10,000 m.s^-1^, then heated at 55°C for 2 min, then cooled on ice for 2 min, then shaken again, then heated at 70°C for 2 min and cooled again on ice. Suspensions were then centrifuged at 22,000 g, 20°C for 30 min, and the aqueous phases were transferred into sterile 2 mL Phase Lock Gel tubes (Eppendorf, Montesson, France). Then 2 μL of 20 mg.mL^-1^ ribonuclease A in 0.04% EDTA dihydrate and 1% Tris-HCL 1M were added, and heated at 37°C for 30 min. Purification was done using Genomic DNA Clean & Concentrator-10 kit (Ozyme, Saint-Cyr-l'École, France) following the manufacturer's instructions. DNA concentration and purity were measured using Qubit (Thermo Fisher Scientific, Les Ulis, France) and by UV spectrophotometry (NanoDrop ND-1000, Labtech International, Ringmer).

The V3-V4 region of the 16S rDNA gene, nearly 460 bp long, was amplified using the primers F343 5’-CTT TCC CTA CAC GAC GCT CTT CCG ATC TAC GGR AGG CWG CAG-3’ (adapted from Liu *et al.*, 2007) and R784 5’-GGA GTT CAG ACG TGT GCT CTT CCG ATC TTA CCA GGG TAT CTA ATC CT-3’ (Andersson *et al.*, 2008) as previously described (Lazuka *et al.*, 2015). The PCR assay conditions consisted of an initial denaturation step at 94°C for 60 sec, followed by 30 cycles of denaturation at 94°C for 60 sec, annealing at 65°C for 60 sec, and extension at 72°C for 60 sec, and a final extension at 72°C for 10 min. The PCR mixture contained 1 μL of a 10mM deoxynucleotide triphosphate mixture (GE Healthcare, Buc, France), 25 μM of each primer (Eurogentec, Angers, France), 5 μL of 10X PCR buffer, 2.5 U Taq DNA polymerase and 10 ng of metagenomic DNA. All dilutions were made under sterile conditions with water BPC grade and all products were purchased from Sigma–Aldrich (Saint Quentin Fallavier, France) unless specified otherwise. Amplicons were sequenced using Illumina Miseq technology. The average depth of the 71 samples (36 at 9 and 20 days of ripening minus one lost) is 28539 reads. Amplicon data from high-throughput sequencing were analysed using Find Rapidly OTUs with Galaxy Solution (FROGS), v3.0 (Escudié *et al.*, 2018). First, 250 bp Illumina reads were merged using VSEARCH tool (Rognes 2016) with expected amplicon size between 400 and 480 bp and a 0.02 mismatch rate. Next, bacterial 16S rDNA gene sequences were clustered into operational taxonomic units (OTUs) using Swarm with a threshold of 97% pairwise identity and a denoising step (Mahé *et al.*, 2014). Then, chimeras were detected *denovo* and removed from the database using VSEARCH tool (Rognes *et al.*, 2016) with UCHIME algorithm (Edgar *et al.*, 2011). We kept only OTUs representing more than 0.005% of total reads, as well as OTUs present in at least three samples. For each OTU, taxonomic assignment was determined with Silva-132 ([https://www.arb-silva.de/](https://www.arb-silva.de/documentation/release-132/)) and 16S rDNA RefSeq databases (<https://blast.ncbi.nlm.nih.gov/Blast.cgi>). The OTUs were then grouped into genera, and the annotations of the two databases were merged. In case of discrepancy, the blast annotation was chosen only if the bootstrap value was greater than 97%, otherwise the Silva annotation was chosen. Only genera representing at least 1% of relative abundance were retained for analyses. Four usual diversity parameters have been calculated on OTU compositions: Shannon index, Simpson index and Bray-Curtis dissimilarity. Formulas are available in Supplementary Table 1C.

4. **Percentage of cheese slices covered by blue**

We estimated the area of cheese covered by blue on fresh inner cheese slices, which depends on the formation of cavities in cheeses, the growth of *P. roqueforti* within cavities and its sporulation ability, as the blue color is produced by melanin in *P. roqueforti* spores. For this, we cut all 20, 90 and 180-day cheeses in half and took three pictures of each fresh slice with a Canon PowerShot SX410 IS (JPEG format, 5152x3864 pixels, 100 ISO speed, without flash). We analysed pictures using imageJ 1.52n (Schneider *et al.*, 2012; Figure S2): (i) brightness and contrast of raw pictures were standardized with stack contrast adjustment plugin with a reference picture (Figure S2A,B) (ii) using a rectangle selection, pictures were manually cropped to only keep cheese slices (Figure S2C), (iii) the red channel was segmented using a grayscale threshold of 102 to discriminate colonized cavities from inner white part or empty cavities (Figure S2D), (iv) the number of dark pixels in the cavities was counted and divided by the total number of pixels of cheese slices to obtain the value of the percentage of blue area in the slice. Within dark patches in the cheese cavities colonized by the fungus, all pixels were considered dark (script at <https://gitlab.com/snippets/1945218>).

5. **Biochemical analysis**
   a. **Physicochemistry**

We performed standard physicochemical measurements on the cheeses: we measured dry and fat over dry matter content in 9, 20, 90 and 180-day cheeses according to the reference methods NF EN ISO 5534 and NF V04 287B, respectively (see acronyms below). Moisture content of the defatted cheese (MDC) was calculated at the same stages following the reference method IDF 1982. We measured total, soluble and non-protein nitrogen contents in 20, 90 and 180-day cheeses according to the reference methods ISO 8968-1, FIL 224 and ISO 8968-4, respectively. Chloride concentrations were measured on 90 and 180-day cheeses following NF EN ISO 5943 (2007) and salt content was calculated from chloride molar mass following IDF 1988. Water activity (Aw) was determined in agreement with the standard NF EN method ISO 21807 (ISO 2004) at 25°C, with a LabMaster (Novasina, Lachen, Switzerland), on 90 and 180-day cheeses (NF: normalisation française; EN: european committee for normalization, https://www.cen.eu/; ISO: international organization for standardization, https://www.iso.org/; AFNOR: association française de normalisation, <https://www.afnor.org/en/>; IDF: international dairy federation, FIL: fédération internationale du lait, <https://www.fil-idf.org/>). Cheese core pH was directly measured using a laboratory pH-meter (CG 841, Schott, Mainz, Germany) with an Ingold electrode 406 MX (Mettler Toledo S.A, Viroflay, France), on 9, 20, 90 and 180-day cheeses.

We measured glucose, lactose, lactate, acetate and butyrate in 9 and 20-day cheeses by high performance liquid chromatography (HPLC) following Pham 2016. Using a scalpel, we cut and weighed 10 g of cheese and mixed it in 90 mL of NaCl-peptone solution (0.85% NaCl, 0.1% Pepton). The mixture was homogenized with a Stomacher for 5 minutes and allowed to stand for 10 minutes; 30 mL of the cheese homogenate were placed in a 50 mL Falcon tube (Eppendorf, Montesson, France) and incubated for 60 minutes at 40°C. The sample was then centrifuged for 30 minutes at 8,000 g (Beckman Avanti J-25I, California, USA). After centrifugation, the samples were placed on ice for one hour in a 50 mL tube. The solid grease block was then pierced by means of a spatula and the liquid was filtered through a 0.45 µm filter. This process removes most solidified milk fat. The filtrate was heated in a water bath at 25°C. The pH of the heated filtrate was adjusted to 4.6 with 10% phosphoric acid to precipitate the water-soluble casein. The acidified supernatant was centrifuged for 30 minutes at 8,000 g in Falcon tubes. After centrifugation, the acidified supernatant was filtered through a 0.45 μm filter (Infochroma AG, Zug, Switzerland). The samples were frozen for two hours, which precipitates the small particles of the soluble casein and the solidified fat. The cheese extract was thawed and filtered through a 0.45 μm filter in order to separate the adhered caseins and the fat particles. Fermentation supernatants were centrifuged for 15 min at 22,000 g, then transferred into a new reaction tube and diluted when required with MilliQ water. The diluted samples were filtered again through a 0.45 μm filter into a 1.5 mL Crimp Neck Vial (BGB Analytik AG, Böckten, Switzerland) immediately closed with a crimp cap for determination of short-chain fatty acids in fermentation supernatants by high performance liquid chromatography (HPLC Accela, Thermo Fisher Scientific, Les Ulis, France). Metabolites were separated on an ion exchange column Rezex ROA-Organic Acid H+ (300 x 7.8 mm, Phenomenex, Paris, France) with an Erosa Nr. 1 needle (Rose GmbH, Barkhausen, Germany) and a Injekt 2 mL syringe (Braun, Minden, Germany). The pressure at the beginning of the gradient was 103 bar and the column temperature 40°C. The flow was 0.4 mL.min^-1^ and the solvent was 10 mM sulphuric acid (Merck, Massachusetts, USA) in MiliQ water. The injection volume was 20 µL and one analysis took 90 minutes. All products were purchased from Sigma–Aldrich (Saint Quentin Fallavier, France) unless specified otherwise. Data were identified by refraction index detection and quantified (g.L^-1^) using ChromQuest (Thermo Fisher Scientific, Accela, Wohlen, Switzerland) based on a linear regression according to standard solution.

b. **Metabolite extractions and analyses**

We tested whether the four populations displayed different proteolytic and lipolytic activities. For this goal, we analysed 90-day cheeses, following two procedures of extraction (water or organic solvents) and UHPLC-MS analysis. Using UHPLC-MS analysis on water extracts of the cheeses, we analyzed whether the *P. roqueforti* population influenced the quantities of water-soluble compounds. Water-soluble metabolites were extracted using the method previously described (Le Boucher *et al.*, 2013). Briefly, 90-day cheeses were frozen at -80C° until extraction. Cheese samples of 10 g were fivefold (w/w) diluted in deionized water (Milli-Q Reagent Grade Water System, Millipore Corporation, Billerica, Massachusetts, USA). The mixture was then blended for 1 min and the homogenate was centrifuged at 10,000 g for 10 min. The supernatant was further centrifuged on centrifugal filter units, with 10 kDa cutoff (Vivaspin 20, Sartorius, Palaiseau, France) at 8000 g for 30 min. The filtrate was stored at -80C° until analysis. After thawing, the samples were filtered again (0.22 µm) and diluted when required in Formic acid 0.1% for analyses by ultra-high performance liquid chromatography mass spectrometry (UHPLC-MS; UHPLC Ultimate 3000 and HR-MS-Q exactive, Thermo Fisher Scientific, Les Ulis, France).

For UHPLC, metabolites were separated on Hypersil Gold phenyl (Length = 15 mm, Internal diameter = 2.1 mm, Particles size = 3 µm, Thermo Fisher Scientific, Les Ulis, France). The pressure at the beginning of the gradient was 120 bar and the column temperature 25°C. The flow was 0.25 mL.min^-1^ and the solvent were HPLC-quality acetonitrile (D) and ultra-pure water plus nonafluoro pentanoic acid (3 mM) (B). The elution gradient was as follows: 4 min at 98% B + 2% D, then 98% to 2% B in D for 6 min, level at 2% B and 98% D during 3 min. The injection volume was 5 µL and the injector temperature 7°C. The duration of one analysis was 14 minutes.

Mass spectrometric detection was performed with a Hybrid Quadrupole-Orbitrap with a heated electrospray source operated in the positive ionization mode. Full scans of mass ratio were acquired with a scan range of 3.7 scan.sec^-1^ and a mass range from 50-700 Da with a resolution of 70,000.

To check if the extracted and analyzed water-soluble compounds (mass range from 50-700) were mostly amino-acids and small peptides, the data were combined into a single matrix by aligning peaks with the same mass-retention time pair together and annotated using a spectral database as previously described (Hébert *et al.*, 2013). As more than 90% of the total integrated surface area is composed of amino acids, this metabolic analysis represents an excellent proxy for proteolysis (Figure S3B).

In order to analyze whether the *P. roqueforti* population influenced the quantities of free fatty acids and residual glycerides in 90-day cheeses, we develop a new extraction procedure. We coupled a global extraction (accelerated solvent extraction with Hexane-Isobutanol) with a UHPLC-MS analysis in positive (triglycerides) and negative (fatty acids) ionization modes. All the steps were optimized using standards of fatty acids and triglycerides, even though the commercial triglycerides used are not found naturally in milk.

For the accelerated solvent extraction (ASE 350, Dionex, USA), 0.5 g of each cheese were precisely weighed and mixed with 1 g of diatomaceous earth as dispersant and desiccant to improve extraction performance. The mix was introduced into a stainless steel cell (5 mL) to which we added 200µL of heneicosanoic acid (C21:0 à 25 mg.mL^-1^ in Hexane/isopropanol 3:2 v/v; Fluka Sigma-Aldrich) as extraction standard. Three extraction cycles using Isopropanol: Hexane (2:3 v/v) were performed. Three static cycles of 15 min were applied. The extraction solvent was pumped through the extraction cell fitted with a cellulose filter and a stainless steel frit. Each stainless steel cell was maintained at a pressure of 100 bar and a temperature of 110°C throughout the extraction. All resulting extracts of a sample were automatically combined in the same vial (15 mL) and stored at -20°C until analysis. After thawing, the sample were filtered again (0.22 µm) and eventually diluted when required in Isopropanol: Hexane (2:3 v/v) by UHPLC-MS (UHPLC Ultimate 3000 and HR-MS-Q exactive, Thermo Fisher Scientific, Les Ulis, France).

For UHPLC, metabolites were separated on C18 Hypersil Gold column (50 mm x 2.1 mm ID, 1.9 µm; Thermo Fisher Scientific, Les Ulis, France). The pressure at the beginning of the gradient was 120 bar and the column temperature 25°C. The flow was 0.25 mL.min^-1^ and the solvent was water/acetonitrile 60/40 v/v with 0.1% formic acid and 10mM ammonium formate (A) and isopropanol/acetonitrile 90/10 v/v with 0.1% formic acid and 10 mM ammonium formate (B). The elution gradient was as described in supplementary data (Supplementary Table 4A). The injection volume was 5 µL and the injector temperature 7°C. The duration of one analysis was 25 min.

Mass spectrometric detection was performed with a Hybrid Quadrupole-Orbitrap and a heated electrospray source operated in the positive and negative ionization mode. Full scans were acquired with a scan range of 3.7 scan.sec^-1^ and a mass range from 70-1000 Da with a mass ratio resolution of 70,000. To validate the analytical repeatability and quality, a pool of all the samples was injected at the beginning and the end, as well as randomly all along the analytical run.

Non-targeted analysis has been done using R software v3.5.1 (<http://www.r-project.org/>) with “MSnbase” v2.8.3, “xcms” v3.4.2 and “CAMERA” v1.38.1 packages (Gatto *et al.*, 2012, Smith *et al.*, 2006, Kuhl *et al.*, 2012, respectively) (script at <https://gitlab.com/snippets/1945216>).

For quantification, the data were quantified (ng.g^-1^ wet weight) using TraceFinder software v3.3 based on a linear regression according to the calibration solutions (Supplementary Table 4B). For triglycerides, as the product ions are NH_4_ adducts [M+NH_4_]+, we were able to identify by their precise mass 19 potential triglycerides (Supplementary Table 4C).

c. **Volatile compound extraction and analyses**

We investigated the identity and abundance of volatile compounds involved in flavor and aromas. The volatile compound extraction was performed using a dynamic headspace (DHS) system with a Gerstel MPS autosampler (Mülheim an der Ruhr, Germany). The sorbent material used was Tenax TA (2,6-diphenylene oxide polymer, Gerstel) conditioned before use as recommended by the manufacturer.

Three grams of cheese sample, previously crushed and stored at -80°C, were weighed precisely into a 20 mL glass vial without thawing. The samples were then stored at 4°C for 16 hours before analysis in order to stabilize the interactions between matrix and volatile compounds. The samples were then transferred to the sampling support (10°C) and analysed randomly. First, the sample was thermostated at 40°C for 3 min and stirred (28 g). Subsequently, the sample headspace was purged with a helium flow of 30 mL.min^-1^ for a total purge volume of 300 mL and the analytes collected at 30°C. In order to reduce the amount of aqueous vapour sampled, the tube was dried at 30°C with a H_e_ flow of 50 mL.min^-1^ with a total volume of 300 mL.

Thermal desorption and cryofocusing were performed using a Thermo desorption unit (TDU) and a cooling injection system (CIS), respectively. The desorption of aroma metabolites was carried out in a solvent venting mode with the following heating program: from 30° to 290°C at 60°C.min^-1^, with a final hold time of 7 min. Analytes were then cryofocused in the CIS injector cooled at -100°C and successively desorbed to 270°C at 12°C.sec^-1^, with a final holding time of 5 min. The temperature of the transfer line between the TDU and the CIS was kept constant at 300°C.

Gas chromatography-mass spectrometry analysis was performed with a 7890B Agilent GC system coupled to a quadrupole mass spectrometer Agilent 5977B (Santa Clara, United States). A HP-Innovax capillary column was used (60 m × 0.32 mm × 0.25 μm film thickness, PEG, Agilent). Injection was performed in a splitless mode (1 min). The carrier gas was H_e_, at a constant flow of 1.6 mL.min^-1^. The column oven temperature program was as follows: initial temperature of 40°C for 5 min, then raised to 155°C at 4°C.min^-1^, and then to 250°C at 20°C.min^-1^ and held for 5 min. The interface was kept at 250°C and the ionization mode was electron-impact (70 eV). All data were recorded using the software Agilent Mass Hunter Qualitative Analysis B.07.00. Aroma metabolites were identified by NIST 17 spectral library. Volatile quantification was performed using signal abundance (TIC - Total Ion Current).

4. **Statistical analyses**

For statistical tests of microbiological counts, image analysis, physico-chemistry analyses, the following model was applied:

$$Y=Population+Stage+Population*Stage+\left( 1\vee Assay \right)$$

Where Y is the dependant variable, Population the population as a fixed effect, Stage the maturation day as a fixed effect and (1|Assay) the assay as a random effect. For statistical test of metabolomic analyses, we applied the same model without any stage effect (only 90-day cheeses were analysed). For the metabarcoding analysis, as the composition data is poorly compatible with mixed model we applied:

$$Y=Population+Stage+Population:Assay$$

All models and possible data transformations are specified in the statistical tables. All statistical analyses were performed with the R software (<http://www.r-project.org/>) using (i) the “lme4” package v3.1.142 (Bates *et al.*, 2015) for the linear mixed models of microbial counts, physico-chemistry and image analysis (ii) the “car” package v3.0.3 (Fox *et al.*, 2019) for analyses of variance on physicochemistry, microbial counts and multivariate analysis of variance on OTU abundance, (iii) the “compositions” package v1.40.2 (Van Den Boogaart *et al.*, 2018) for isometric log ratio transformation of OTU abundances, (iv) the “pls” package v2.7.0 (Bjørn-Helge *et al.*, 2018) for orthogonal signal correction partial least square discriminant analysis (O-PLS-DA) followed by t-tests on jackknife resampling of PLS regression coefficients on metabolic and volatile compounds, (v) the “emmeans” package v1.3.2 (Searle *et al.*, 1980) for post-hoc analyses with estimated marginal means, (vi) the “vegan” package v2.5.6 (Oksanen *et al.*, 2019) for the Bray-Curtis dissimilarity calculation, (vii) the following packages for plots: “ggplot2” package v3.1.0 (Villanueva *et al.*, 2016), “cowplot” v0.9.4 (Wilke, 2019), “gridExtra” v2.3 (Auguie, 2017), “plyr” v1.8.1 (Wickham, 2011), “Cairo” v1.5.10 (Urbanek *et al.*, 2019), and “TeachingDemos” v2.10 (Snow, 2016). For O-PLS-DA, the number of components and orthogonal signal corrections was determined using a minimum root mean square error of prediction (RMSEP) with at least two components (see statistical tables).
